# Primary cilia in the growing limb are preferentially orientated, uncoupled from centriolar position

**DOI:** 10.64898/2025.12.19.695513

**Authors:** T Johnson, S Midha, AM Julé, G Randall, K Apolinova, J Miotla-Zarebska, SN Sansom, TL Vincent, AKT Wann

## Abstract

How cells and their organelles are positioned in three-dimensional, organ level, anatomical context, is rarely investigated. Here we focus on cells, centrioles and primary cilia in the growing limb. Through the cilium’s mechanobiological role in skeletal development, we explored the mechanobiology of morphogenesis.

A transgenic mouse model (Centrin 2-GFP.ARL13B-mCherry), combined with an image analysis pipeline, can map cellular size, positions and orientations, centriole position and ciliary axoneme orientation, all with respect to the anatomy of the epiphysis or growth plate. The line was crossed with an *ift88*^fl/fl^CreER^T2^ line to enable ciliary *ift88* deletion. We used limb immobilization, to test for a role of mechanical forces associated with ambulatory loading, in the organization of these elements and transcriptomics to understand the role of forces in regulating growth plate morphogenic programs.

The pipeline can accurately quantify expected patterns of cell orientation and size through zones of the growth plate. Analysis across thousands of cells, through regions and zones of multiple murine growth plates, reveals cilia prevalence is increased in the periphery, highest in the resting zone in the outer limb, harboring stem cells. Cilia length is greatest in the hypertrophic cells about to die or transdifferentiate, as part of the formation of bone from cartilage by endochondral ossification. The inducible and cartilage-specific, deletion of ciliary gene *ift88,* alters cell orientation and sizes and reduces ciliation in the areas where endochondral ossification is most disrupted, the periphery and expanded hypertrophic zones, linking changes in structure to function. Most strikingly, centriole position, including that of the basal body, from which the ciliary axoneme is extended, is not preferentially organised. In contrast, cilia axoneme orientation is preferentially organised. Axonemes are directed posterior or anterior, 45 degrees to the axis of the limb, irrespective of their position, which is defined by basal body position. Immobilization of the limb for 2 weeks markedly alters the transcriptomic profile of the growth plate, with changes to size and orientation of cells and alterations in matrix and cytoskeletal profiles. Within altered genes, primary cilia genes themselves are regulated, including those indicative of altered cilia signaling such as hedgehog signaling. However, despite the role of cilia in mechanobiology of the growing limb, and ciliary signature within changes to loading of the limb, cilia orientation is unaltered by the removal of ambulatory associated forces.

Patterns of ciliation in control and IFT88cKO mice help reconcile the previously observed anisotropic effects of cilia perturbation, focusing study on stem cell-resting chondrocytes and hypertrophy, when considering the mechanobiological role of cilia in limb development. Endochondral ossification is apparently highly sensitive to ambulatory loading at transcriptomic level, including effects on ciliary genes and signaling. A highly organized orientation of these putative antennae is governed by centriole position-independent mechanisms and is independent to changes to ambulatory loading, indicating a cell intrinsic mechanism.

The resilient position of axonemes in the limb, points to mechano-regulatory mechanisms for how cilia integrate biophysical signals. We propose that predominant ventral or dorsal orientation at 45 degrees to horizonal plane but never parallel to cranial-chordal or medial-lateral axis, ensures multiple signal integration and avoids single signal ‘blindness’.

## INTRODUCTION

Relative cell and organelle position or polarization is essential in many tissue types during morphogenesis, and to maintain homeostasis. Modern imaging preparations and technologies are enabling us to explore these *in situ,* to understand tissue and cellular anisotropy at organ level. Why, for example, some cells, in some locations, respond differentially to others, is likely to be a composite effect of intrinsic genotype and gradients of a range of stimuli and factors that encompasses the biological and physico-chemical. How cells are positioned to sense these, is also critical.

At the turn into the 21^st^ century it became clear that the once thought vestigial, primary cilium, was critical to a plethora of cell and tissue biology. Imaging has been a mainstay during the growth of primary cilia research; the structure and relative orientation of primary cilia has often shaped proposals for the mechanisms underlying ciliary influence[1]. Non-motile and only one per cell, these microtubule-based organelles are often coined as a cellular antennae [2]. In some cell types they project out into the extracellular space, apparently well-placed to sense biological and physico-chemical cues. Despite an appreciation of the importance of these ‘antennae’ to development, health and disease, to quote from a review on this subject [3] “for most cells of the vertebrate body we do not yet understand where the antennae is positioned and where it is pointing”. This has largely not changed in the 14 years since this was written, despite the ever-increasing repertoire and power of microscopic techniques. It’s important to note that in many tissues this “antennae” is not always projected into the extracellular space but embedded in an invagination in the membrane called the ciliary pocket [4]. With the development of scalable imaging pipelines, the mapping of cilia position and orientation, in the tissue context, has now become possible [5, 6] at an organ level.

Cilia are a hub for a plethora of signalling [7], recruiting receptors for many signals, ranging from growth factors [8–10] to mechanical forces [11–15]. However, many mechanistic questions remain, especially in the domain of quite how cilia are positioned to transduce mechanical forces to adapt cell and tissue behavior. The role of putative mechanosensors such as the polycystins or role of the ciliary axoneme itself, remains unclear. Recent elegant studies from the node [13, 14] continue to shed more light on how cilia might integrate biological cues such as bone morphogenetic protein (BMP), fibroblast growth factor (FGF), sonic hedgehog (Shh) signalling, with mechanical cues such as fluid flow [16, 17] helping cells decipher left from right. In this context clearly the position of ‘sensory’ ciliated cells in the tissue, and position and orientation of the ciliary axoneme on the cells, is fundamental to their role in chemo-mechnosensing. Whilst this might also hold true in tissues such as the kidney [18, 19] or the skeleton [20], where pathological changes that define the human ciliopathies are associated with the primary cilia dysfunction [21], descriptions in these tissue contexts remain relatively limited.

Whilst exploring the influence of primary cilia in the post-natal limb, our own work identified an unexpected role for cilia in the mouse growth plate. Here they apparently act to integrate mechanical forces during skeletal growth[22], in a paradoxical manner. In the appendicular skeleton, the growth plate is a cartilage template for bone growth. Within, a complex cascade of cell differentiation events and tissue changes underpin a rapid transition from stem cell, through chondrocyte, to bone, defining endochondral ossification EO[23]. Upon deleting the ciliary gene encoding *Intraflagellar transport protein 88* (IFT88), in chondrocytes, during post-natal growth, we observed an elongation of the growth plate indicative of a failure to mineralize [22] and an expansion of hypertrophic chondrocytes. Appendicular growth plates comprise a stem cell niche [23, 24], which feeds clonal populations organized in the post-natal limb in proximal-distal columns [25] spanning the chondrocytic lineage from resting, through proliferative to hypertrophic, and which ultimately transdifferentiate to become bone in a highly controlled manner [26]. Cilia [27–31] and ciliary mediated signalling including Indian hedgehog (Ihh) [32–35] are established central players in growth plate dynamics, thus disruption was not unexpected, but most mutations or pertubations to either are associated with GP shortening and accelerated growth plate closure. Our work indicated that the effects of IFT88 deletion on EO in adolescence are mechano-dependent. This implied that cilia protected GP dynamics from ambulatory, physiological loading [36]. In particular, the data suggested this role may be more important in the periphery of the limb, which motivated us to map ciliation within the murine adolescent growth plate. Primary cilia orientation has been described in articular cartilage, where it is thought to be influenced by weight-bearing [37] and, in a limited fashion, in the growth plate where consensus is that orientation is preferential to the axis of the limb [3, 37–40]

Given we observed ciliary perturbation had a preferential effect in the growth plate periphery [22], in a mechano-dependent manner, our hypothesis was that ciliary prevalence, length or perhaps position within the tissue would vary between the apparently unaffected central region of the growth plate and the periphery. These regions are predicted to experience differential mechanical stress and likely cell strains due to the geometry of the epiphysis[41]. To accomplish this, we exploited the transgenic model first used to study embryonic ciliogenesis [42] which enables accurate, large scale, visualization of cilia and centrioles in 3D tissues [6]. Furthermore, we exploited genetic deletion of IFT88 and surgical immobilization to remove ambulatory loading and both of which alter course of endochondral ossification. We characterized the cellular and transcriptomic changes upon immobilization. Finally, we analyzed cilia and centrioles across the altered landscape of the growth plate upon removal of ambulatory loading. We find that primary cilia are assembled by a large proportion of chondrocytes through the different zones of differentiation. We find differences in these between the centre of the limb and the periphery, which may explain relative susceptibility of these regions to IFT88 deletion. Primary cilia orientation is indeed highly non-random, but in contrast to previous findings, our larger scale and unbiased analysis indicate a preferential dorsal-ventral orientation. Strikingly, this preferential orientation is not associated with centriolar, and therefore ciliary, position on the cell and thus appears independent to basal body position. Whilst immobilization radically alters the transcriptomic profile of the growth plate, including a signature associated with primary cilia themselves, and cell size and orientation, preferential ciliary orientation is not affected by the removal of ambulatory loading in the limb for two weeks. This suggests cilia orientation may be too important to be controlled by mechanical forces alone and, we suggest, is critical to the mechano-regulatory integrative role cilia play during developmental dynamics.

## METHODS

Methods used for tissue imaging pipeline are captured largely in recent description [6]

### Animals

All mice were housed in the biomedical services unit at the Kennedy Institute, University of Oxford. Mice were housed no more than 6 single-sex mice per cage, in individually ventilated cages and maintained under 12-hour light/dark conditions at an ambient temperature of 21°C. One mouse line was used for this project: Ift88_fl/fl_;Centrin 2-GFP.ARL13b-mCherry (Ift88 _fl/fl_) [22, 42], the control line for conditional knockout of IFT88 previously described (Ift88 _fl/fl_ are used in sequencing analyses Figure 6&7) [22, 43] but now crossed to Centrin 2-GFP.ARL13b-mCherry and are designated IFT88^CTRL^. For all experiments, apart from double neurectomy (off-loaded & contralateral) where only male mice were used, both sexes were used. The mice were fed a certified mouse diet (Special Diets Services: Rat and Mouse No. 3, pelleted, product code 801030), and water ad libitum. Animal husbandry and experiments were conducted in accordance with the University of Oxford ethical frameworks and under a Project license as granted by the UK Home Office. Mice were euthanised by filling a box containing the mice with CO_2_ up to 80% by volume and maintaining its concentration for 5 minutes. Death was confirmed by performing cervical dislocation. Due to the nature of experiments paired with studies exploring IFT88 functional role (IFT88cKO), tamoxifen was administered to all the mice imaged. Hence all mice are termed IFT88^CTRL^. Tamoxifen was dissolved (90% sunflower oil:10% ethanol) at 20 mg/mL by sonication. 50 μL tamoxifen was administered via intraperitoneal injection to mice at 4 weeks of age, for 3 consecutive days, and tibia collected 2 weeks later. For RNA sequencing analyses caging and also therefore date of collection largely overlapped with sample groups, as confirmed by WGCNA analyses. This was a necessity as for example bedding materials have to be different, for welfare reasons, in immobilised groups, however this lack of cage controlling is a limitation.

### Tissue collection & processing

The hind leg knees were dissected and trimmed of excess fat and muscle tissue. The tissue was fixed (10% neutral buffered formalin, room temp-RT, 4h) then in 2.5% formalin, 4°C, overnight. The tissue was then transferred to 30% sucrose in deionised water (diH2O) and gently rotated (RT,overnight or 4°C, 72 h). The tissue was subsequently cryoembedded in super cryoembedding medium, with the kneecap facing down, and stored at –20°C. 50 μm-thick tissue sections were cut with a cryostat from the centre of the joint/limb, using Kawamoto [44] cryofilm tape. Throughout, care was taken to keep the tissue in the dark to avoid fluorescent signal loss. For phalloidin staining, prior to DAPI staining, sections were stained with Alexa Fluor 568 phalloidin (Invitrogen) for 45 minutes, in the dark, at room temperature. Sections were thawed, washed with three 5-minute washes in phosphate buffered saline (PBS) and nuclei stained with 10 μM 4’,6-diamidino-2-phenylindole (DAPI) for 1 minute. After three 5-minute washes in PBS, the sections were mounted with SlowFade Glass Antifade Mountant (invitrogen) and left at room temperature overnight. Imaging was performed the following day. The same protocol was applied to kidney tissues also imaged.

### Confocal imaging

Images were acquired using a Zeiss 980 Airyscan 2 confocal microscope with an immersion oil 40x/1.4 objective. Images were taken in the middle, lateral and medial regions of the growth plate [6]. 40 μm-thick Z-stacks of 0.15 μm intervals were acquired in the DAPI, mCherry and GFP channels. The superresolution SR-8Y Airyscan mode was used. The resolution in x/y of this mode was 120/160nm and in z 450nm. The images were airyscan processed and stitched using the ZEN software (Zeiss).

### Image analysis pipeline

The arivis Vision4D software (Zeiss) was used for analysis as described [6]. A custom pipeline (Supplementary figure 3) was created which consisted of the following steps; A Cell detection step-using the Cellpose Python Segmenter, cells were detected based on background GFP and DAPI signal [6]. Detected cells outside the volume range of 200 to 30,000 μm_3_ were excluded. Detected cells were stored as objects. Primary cilia and centriole detection step: Primary cilia were detected based on mCherry signal and only included if they were between 0.3 and 12 μm^3^ in volume. Centrioles were detected based on GFP signal and included if between 0.02 and 2 μm^3^ in volume. Pairs of centrioles that were detected together were split in two. Cilia and centrioles were matched to their corresponding stored cells based on overlapping signal. Results export: Features of the cells, primary cilia and centrioles were exported in an Excel spreadsheet. Included features were; number of Children (children of cells are primary cilia and centrioles), name, Parent Names (Parent cell for primary cilia and centrioles), 3D Oriented Bounds (minimum outer box of the object), Short side, Middle side and Long side, Angle XY, Angle XZ and Angle YZ, Bounding Box coordinates X1, X2, Y1, Y2, Z1, Z2, SizeX, SizeY, SizeZ (length between the coordinates), Centre of Geometry X, Y, Z, Object’s First & Last Plane and Count (number of planes it occupies), Sphericity, Volume, Surface Area

### 3D spherical coordinates

Spherical coordinates are a three-dimensional coordinate system used to define the position of a point in space by its distance from a fixed origin and two angles (Figure S1. A-C).

**Supplementary Figure 1:**
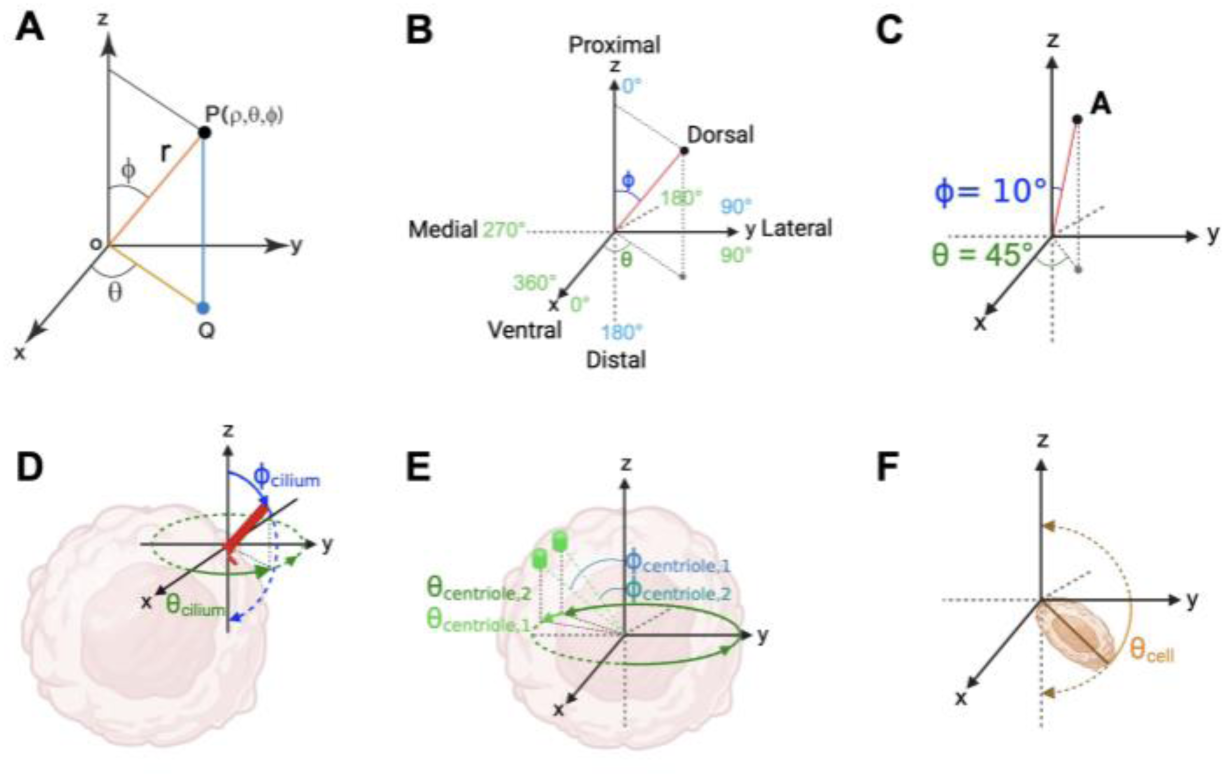
Spherical coordinates. (A) Spherical coordinate system, (B) with anatomical directions and angle values. (C) Example point A has spherical coordinates Φ = 10° & θ = 45°. Spherical coordinate system position to measure primary cilium axoneme orientation (D) and centriole position (E) spherical coordinates, and cell angle θ_cell_ (F).

A point is described using three values; r – the radial distance from the origin, Φ – the polar angle measured as the inclination from the positive z-axis (0→ → Φ → 180→), θ – the azimuthal angle measured as the projection on the xy-plane from the positive x-axis (0→ → θ → 360→). When analysing 3D orientation of primary cilia, r is the length of the primary cilium and its 3D orientation is described by Φ_cilium_ and θ_cilium_) (Figure S1D). These angles have previously been used to describe ciliary axonemal orientation [45, 46]. For the centrioles, the origin of the system is the centre of the cell and Φ_centriole_ and θ_centriole_ describe the position of the centriole near the cell surface (Figure S1E). To measure cell orientation, the θ_cell_ was described as the angle between the major axis of the cell and the y-axis in the yz plane (Figure S1F). An R code pipeline was developed to calculate relevant measures from arivis export. Features calculated were; Cell volume, Cell θ angle to medial-lateral plane (θ_cell_), Primary cilium length, Primary cilium 3D orientation, Primary cilium Φ angle (Φ_cilium_), Primary cilium θ angle (θ_cilium_).

### Micro-computed tomography (MicroCT)

Knee joints were imaged using a Quantum FX Microcomputed Tomograph scanner (Perkin Elmer). Knees imaged kneecap down using the following acquisition parameters: 90 kV voltage, 160 μA current, 5 mm field of view, 3 minutes of acquisition time (fine scanning). Scans (DICOM files) were loaded into ImageJ/Fiji as image sequences. The femoral growth plates were chosen as an anatomical reference to orient the knee joints. The images were then re-sliced to produce coronal sections through the whole joint. The disappearance of the lateral meniscus from view at the distal end of the joint in the image sequence was chosen as an anatomical reference to select the section to measure. The image was converted from greyscale to thresholded by applying a threshold function with the Ostu method with parameters 125 to 255 intensity values. 9 measurements were made across the growth plate in both the greyscale and thresholded images – 3 in each of the medial, middle and lateral regions (Figure S2). The measurements were averaged to calculate growth plate length.

### Histology

Knee joints were fixed (10% formalin,24 h, RT). Joints were decalcified (0.5 M ethylenediaminetetraacetic acid, paraffin embedded, and coronally sectioned at 4 μm thick. Sections were collected from the middle of the joint, cleared in xylene, rehydrated in graded alcohols and stained before brightfield imaging (Zeiss Axioscan 7 10x).

### Sequencing of growth plates – sample collection

Tissue extraction was performed using a dissection microscope. Each limb was dissected at the femur, just above the knee and cleaned thoroughly of any adjacent muscle and soft tissue. Using bent forceps, the limb was held firmly just above and below the growth plate (GP) and snapped towards the diaphyseal end to induce a clean fracture through the growth plate. This resulted in two GP–containing segments: one associated with the femoral end and one with the diaphysis. Under a dissection microscope, the GP, visible as a shiny surface, was gently scraped and collected in RLT buffer, taking care not to penetrate into the underlying bone (for before and after removal see Figure 6A). The tissues were suspended in 350μl BufferRLT (supplied in the kit) with 1:100 β-mercaptoethanol (Sigma-Aldrich, Missouri, USA) and processed for RNA extraction using the RNeasy Micro Kit (Qiagen, Hilden, Germany). Two GPs (i.e. from one mouse) were pooled and homogenised by vortexing for 10 minutes at RT in 1ml TRIzol, followed by repeated passage through syringes with progressively smaller gauge needles to ensure thorough homogenisation of the tissue.

The homogenate was centrifuged at 13,000rpm for 10 minutes at 4°C, and the supernatant was transferred into a fresh tube and mixed with 100ml 1-bromo-3-chloropropane (BCP; Sigma-Aldrich, Missouri, USA). The solution was vortexed until it became milky pink, and centrifuged at 13,000rpm for 20 minutes at 4°C to separate the phases. The resultant aqueous layer was transferred into a fresh tube, mixed with half its volume of 100% ethanol (175μl), and loaded onto RNeasy MinElute spin micro column.

Subsequent RNA extraction was performed as per manufacturer’s protocol with minor modifications. Briefly, following DNase I digestion and washing with Buffer RW1, the column was washed with 500 µl Buffer RPE and centrifuged at 10,000 rpm for 15 seconds. The flow-through was discarded and the column was washed with 500 µl freshly prepared 80% ethanol and centrifuged at 10,000 rpm for 2 minutes. Columns were then centrifuged at 13,000 rpm for 5 minutes with the lids open to remove residual ethanol. RNA was eluted in 14 µl RNase-free water after a 10 minutes incubation at RT.

The extracted RNA was assessed for quality and concentration using NanoDrop spectrophotometry, and RNA integrity was evaluated using RNA Integrity Number measurements, before proceeding with sequencing.

### RNA sequencing analysis

Raw FASTQ files were quality-controlled and aligned to the mouse genome (build mm10) using the txseq pipeline (https://txseq.readthedocs.io/en/latest/index.html). Reads were quantified by calling salmon (version 1.10.0). Salmon counts were imported into R (version 4.3.1, R Foundation for Statistical Computing) and aggregated at the gene level using tximeta (version 1.20.3 [47]. Information from Ensembl version 110 was used for annotations and transcript-to-gene conversion. Low expression genes, defined as those which did not show at least 5 read counts in at least 2 distinct samples across the whole dataset, were excluded from the differential expression analysis (DEA) contrasting WT immobilized to WT naïve mice. The analysis, run in R (version 4.4.0) using DESeq2 (version 1.44.0) [48], thus tested for differential expression of 19659 genes. Log2 fold change (LFC) values estimated by DESeq2 were corrected using the ashr shrinkage estimator (version 2.2-63 [49]), and significance in differential expression was then defined as an adjusted p-value < 0.05 and |LFC| > 1. Geneset enrichment analyses (GSEA) were performed using clusterProfiler (version 4.12.0 [50]). Genes that passed a p-value threshold of < 0.2 were ranked by ashr-corrected log2 fold change, and the gene ontology (GO) database (biological processes subset) was used to test enrichment of key terms. Results were filtered for GO terms of reasonable size (10 to 500 genes, to avoid generic enrichment), and significant enrichment was defined as an adjusted p-value < 0.05 and a normalized enrichment score (absolute value) |NES| > 1.5.

### WGCNA analyses

WGCNA was performed using cornet (link/citation), a wrapper pipeline for the R wgcna package [51]. The input dataset comprised 28 samples was first filtered to include only genes with at least 1 transcript per million (TPM) in at least 3 distinct samples. The data were then normalized using the variance stabilization transformation (vst) from DESeq2. No outlier sample was found at initial QC, and the following parameters were used for further data cleaning: minimum fraction of non-missing samples for a gene to be considered good of 0.5; minimum number of non-missing samples for a gene to be considered good of 4; minimum number of good genes of 5000; minimum number of objects on branch to be considered a cluster of 10. The module detection was run using the stepwise approach. The network type was set to signed-hybrid and with a soft power of 3. Adjacency was characterized using the bicor correlation function, and dist distance function. The TOM type was unsigned. The minimal number of genes to define a module was set to 30, and the dissimilarity threshold before merging modules was set to 0.15. For each module, enrichment in specific GO terms was studied using an over-representation analysis as implemented in the gsFisher package.

## RESULTS

### Large scale, high resolution confocal imaging and associated imaging pipeline can catalogue cilia across and throughout multiple growth plates

As described in our recent protocols paper [6], we combined high resolution imaging of thick sections, with an imaging analysis pipeline, enabling imaging of thousands of cells from 18 different animals growth plates. Sections studied were anterior-ventrally central as identified by menisci in the joint space and included all regions of GP central to periphery and from SOC down into spongiosa (Figure S2A-E) but we focused on the Toluidine Blue/Safranin-O-stained, proteoglycan and collagen rich growth plate and the chondro-osseous transition to bone (Figure S2A/A^i^).

**Supplementary Figure 2.**
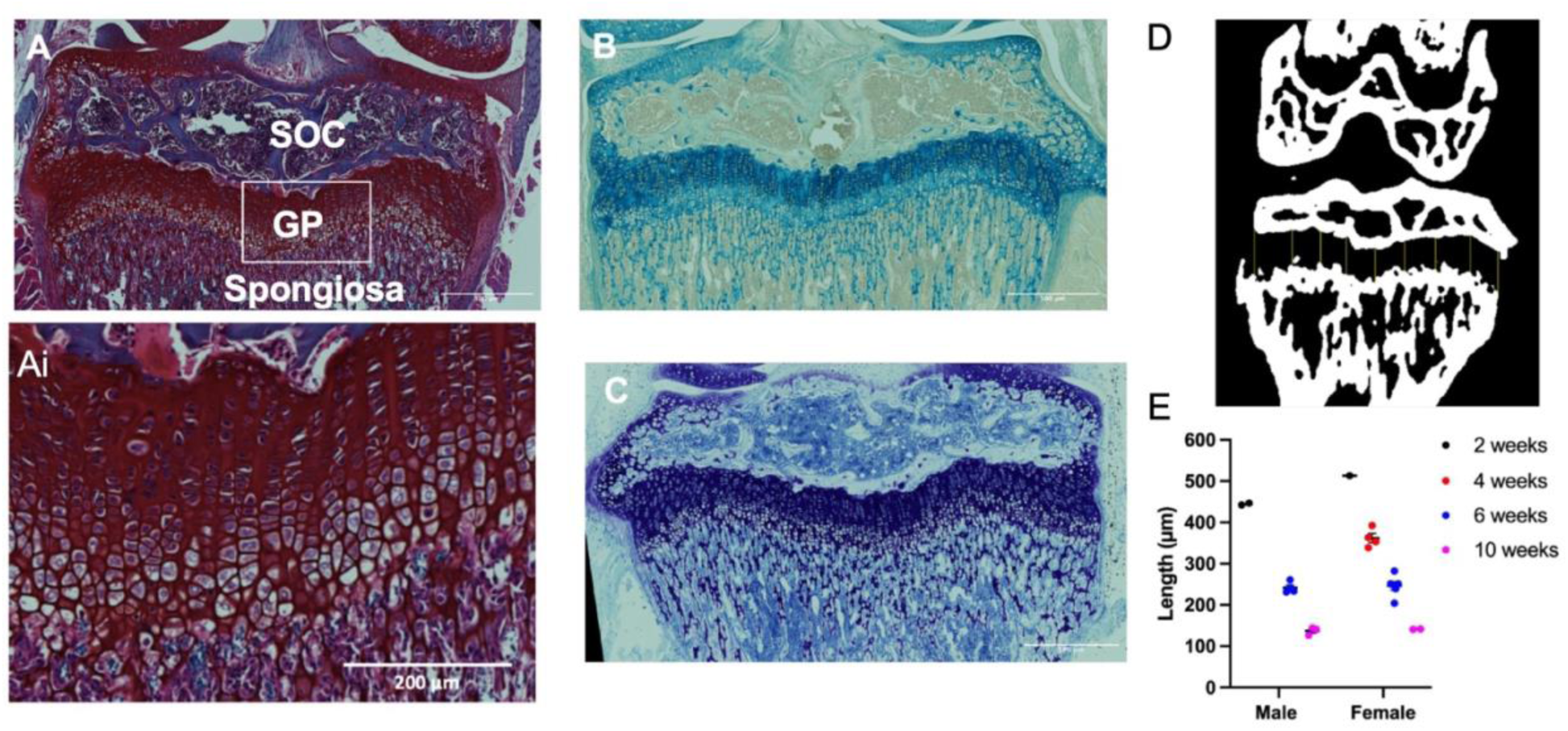
(A) Safranin Orange/Fast green, Alcian blue (B) and Toluidine blue (C) stained sections of tibial epiphysis. uCT image (D) growth plate length measurements (E) Growth plate lengths at different ages. Mean ± SEM. n = 2-5 mice.

Cilia were imaged throughout zones of the growth plate (Figure 1A, resting (B), proliferative (C), hypertrophic (D)). Resolution of 120-160nm x/y, was sufficient to usually discriminate 2 centrioles, one of which is ciliary basal body (Figure 1E/F). Additional staining, including phalloidin staining of actin cytoskeleton, was used to confirm cell detection and associate cytoskeletal structure with morphology. Phalloidin staining of cells within GP showed characteristic hypertrophy within tissue sections (Figure 1G). The pipeline (Figure 1H-I, described in S3) detects cells, cilia and centrioles and maintains their relationships at single cell level, matching objects to a cell. Imaging in all axes, is of sufficient resolution to capture orientation in 3D (Fig1L-N), but requires correction for Z blur later in the analysis (Figure S3).

**Figure 1.**
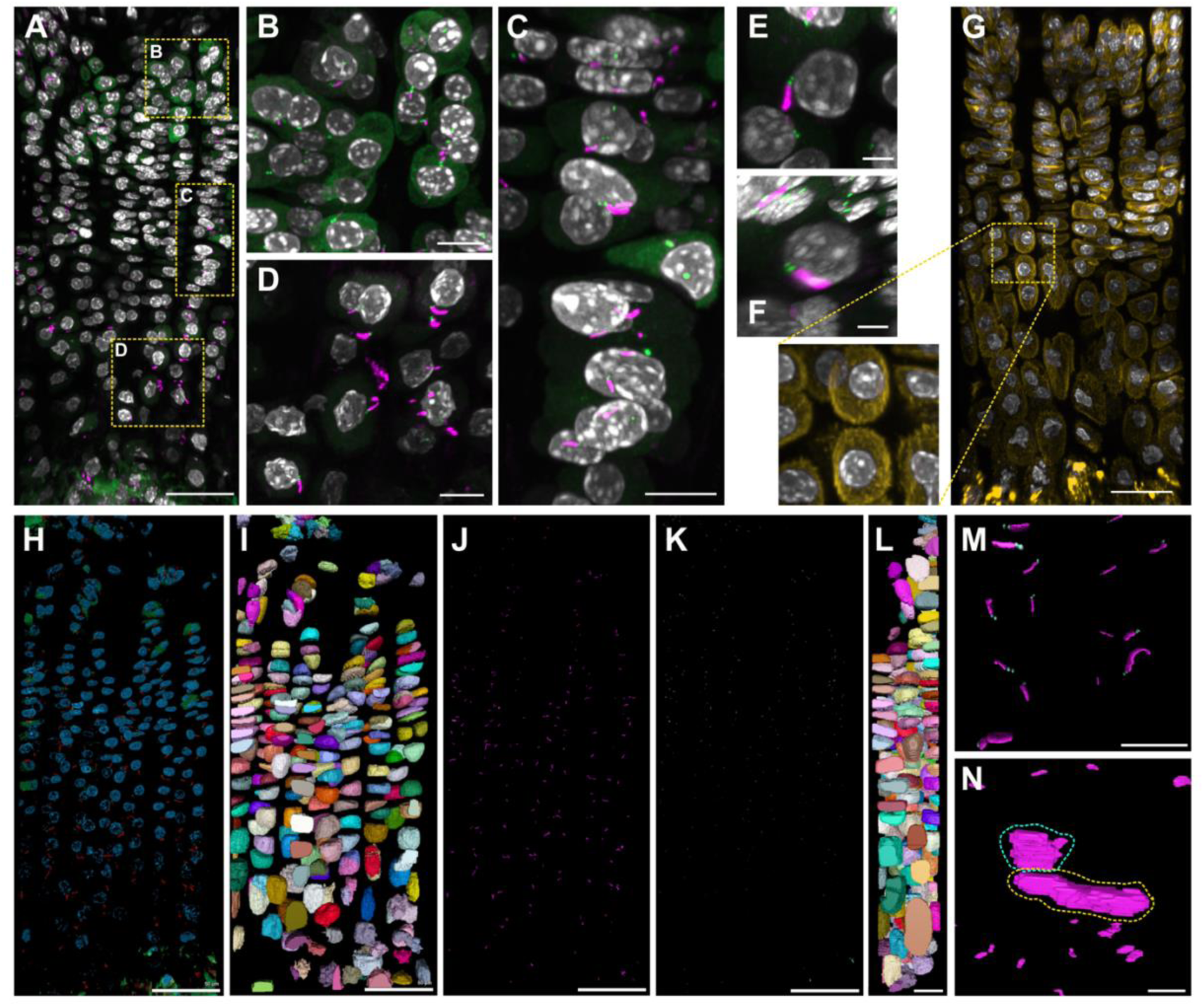
Ciliary Imaging pipeline. (A) Primary cilia in the growth plate a of a 6-week old Ift88^CTRL^ mouse. Scalebar: 40 µm. Yellow boxes indicate regions that are enlarged in the resting (B), proliferating (C) and hypertrophic (D) zones. Scalebars: 10 µm. (E) Single cilium viewed from the front. (F) The same cilium viewed from the side. Scalebars: 2 µm. (G) 6-week old Ift88^CTRL^ murine growth plate stained with DAPI and phalloidin enabling confirmation of cytoplasm and hypertrophy. Scalebar: 40 µm. (H-I) Images show appearance in the software. Cell (I), primary cilia (J) and centriole (K) detection. Scalebars: 50 µm. (L) Side view of the cell detection. Scalebar: 20 µm. (M) Primary cilia and centriole detection together. Scalebar: 20 µm. (N) Zoomed in side view of primary cilia detection. Cyan dashed line: cilium pointing sideways; yellow dashed line: cilium pointing backwards. Scalebar: 2 µm.

**Supplementary Figure 3.**
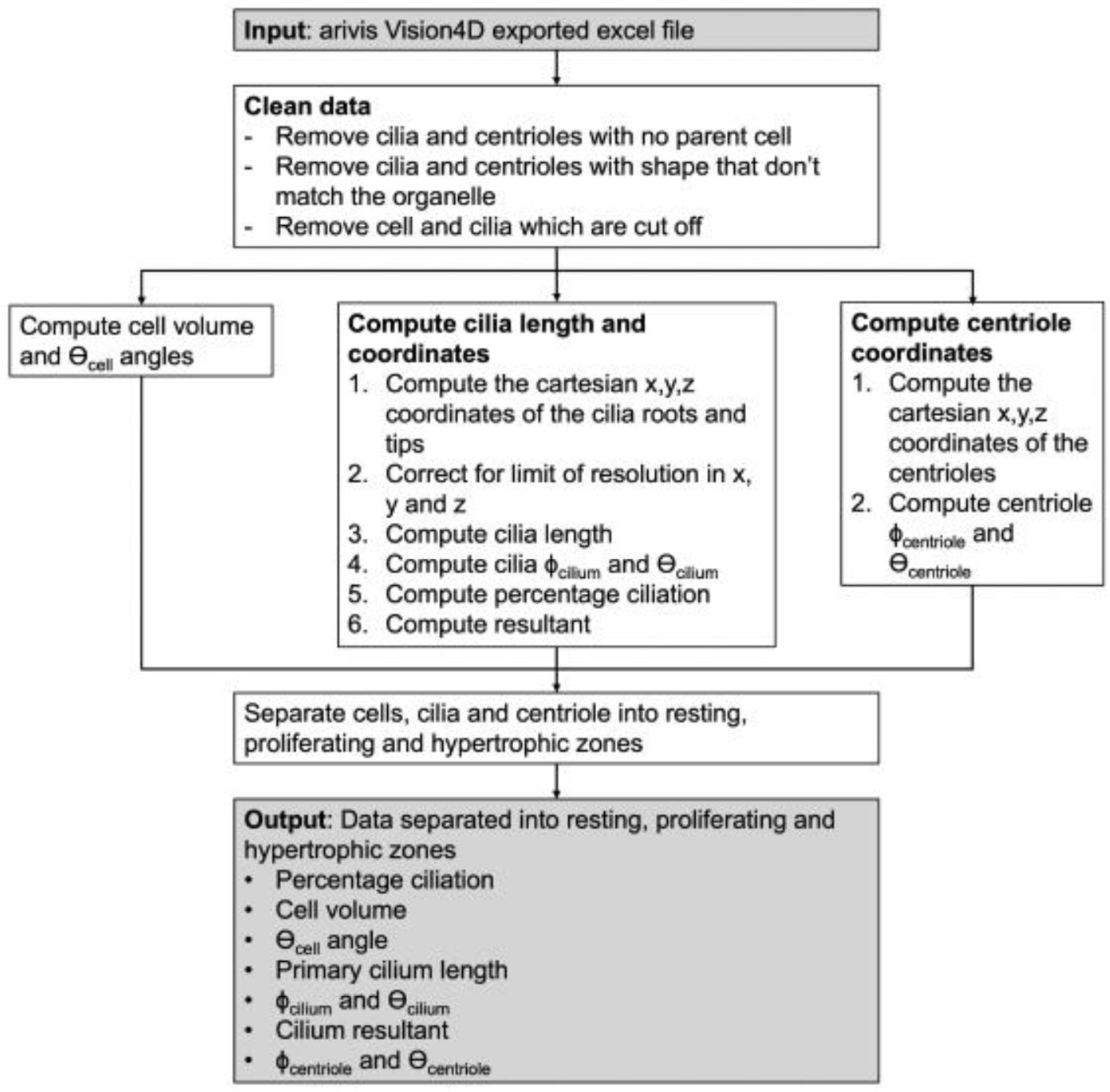
Flowchart describing the steps of the analysis pipeline run in R to calculate cell, primary cilia and centriole characteristics.

### Analysis indicates majority of cells are ciliated, with subtle zonal and regional differences including greater ciliation in limb periphery

Qualitative analysis across GP of mice from 2-10 weeks of age, indicated ciliation was high across all age groups despite increasing senescence of GP and associated shortening of GP length [22] (Figure S4).

**Supplementary Figure 4.**
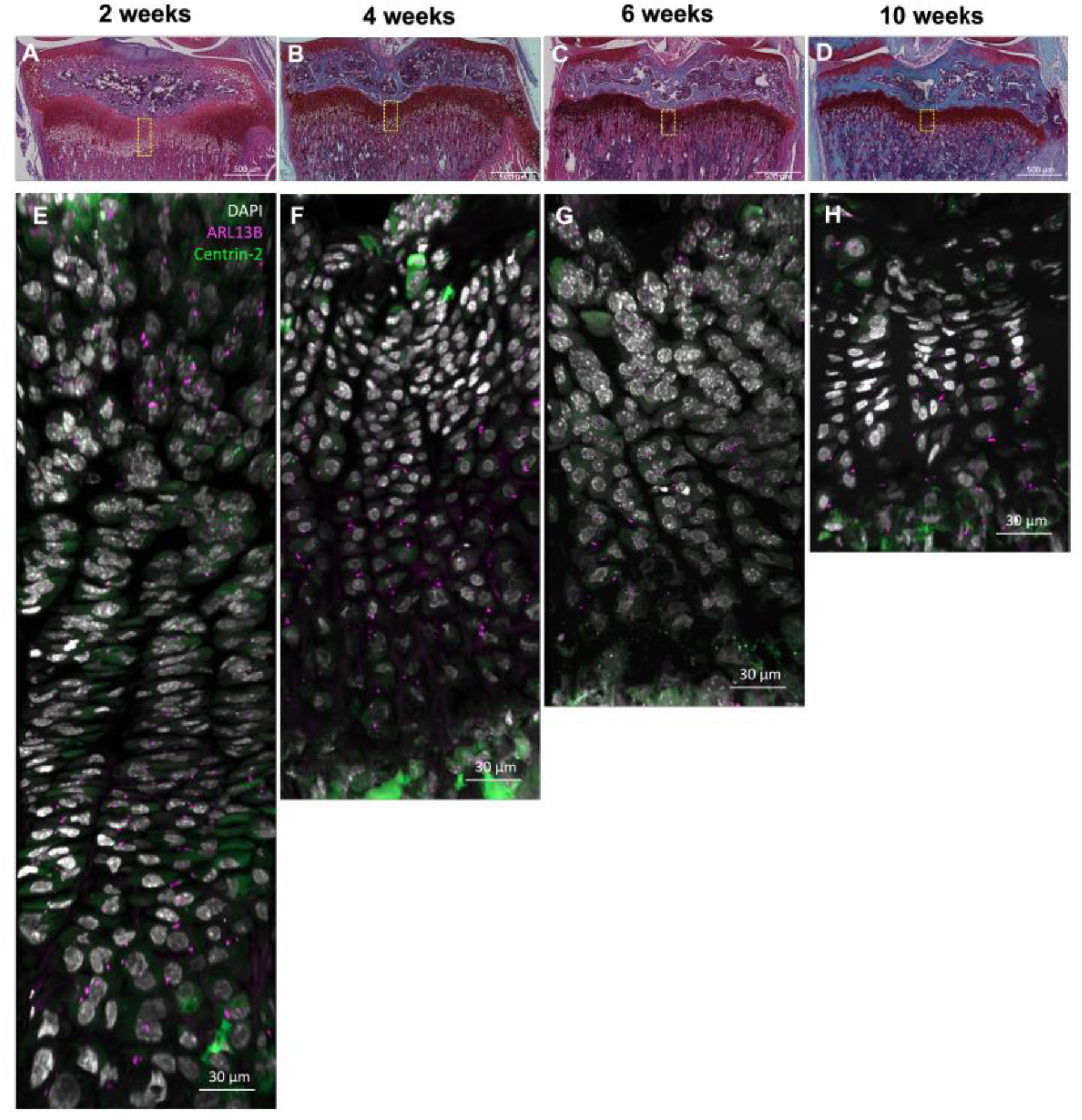
Safranin-O staining. (A-D) and primary cilia imaging (E-H) of murine growth plates at different ages. E, G and H were acquired with the SR8Y Airyscan mode and F with the Co8Y mode (lower resolution, faster). Yellow boxes are representative of the regions of the growth plate imaged for fluorescence.

The previously described, Coveney *et al.* phenotype implied larger, exacerbated effects in hypertrophic zone and periphery of the growth plate[22]. The work implicated cilia in protection from disruptive ambulatory loading, with differential implications in different regions across the GP and, potentially, zones through its length. For example, we hypothesized that the effects being preferentially in peripheral hypertrophic regions was due to increased ciliation in these regions and zone. As such regions (central and lateral) are chosen for analysis (Figure 2A), in order to test this hypothesis and potentially explain the anisotropic phenotype of IFT88cKO [22]. Zones, resting, proliferative and hypertrophic were demarcated manually by cell morphology, (Figure 2B) the pipeline was able to discern the predicted changes in cell size across zones (Figure 2C) and analyze cell orientation in these zones (Figure 2D/E). Analyses quantified the relatively increased horizontal orientation of cells (stacks) in proliferative zone, especially in central region when compared with other zones. The lateral region showed increased heterogeneity in orientation indicative of greater preference of cell orientation in these regions, with respect to central regions, again potentially reflective of differential loading patterns.

**Figure 2.**
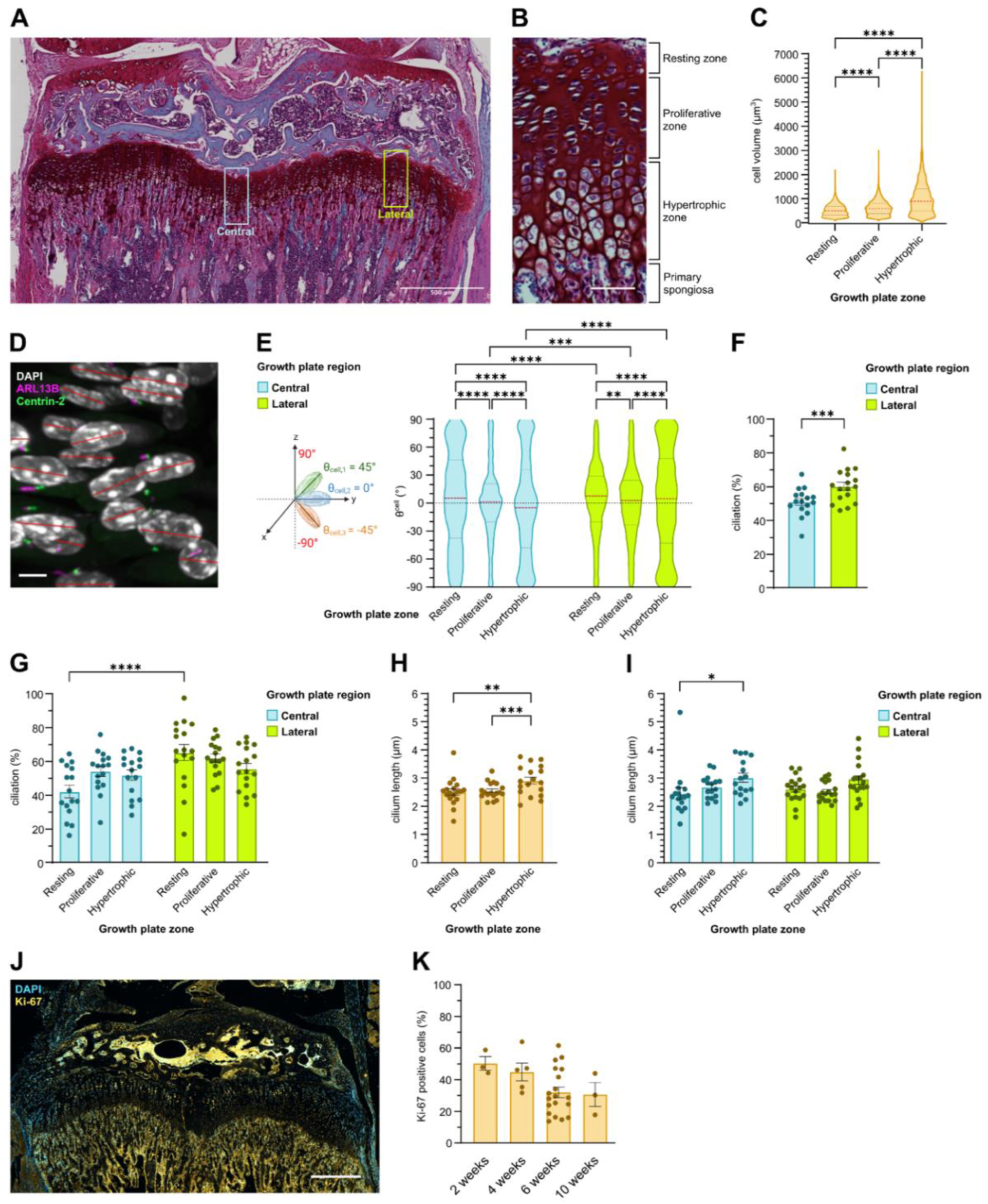
Cell size and orientation, ciliary prevalence and length, across GP zones and comparing middle and lateral regions. Cell organisation differs by zone and region in the 6-week old Ift88^CTRL^ growth plate. (A) Safranin-O stain of the growth plate of a 6-week old Ift88^CTRL^ mouse. Boxes indicate regions that were imaged with fluorescence microscopy. Scalebar: 500 µm. (B) Zoom of histology showing zones. Scalebar: 50 µm. (C) Cell volume in the resting, proliferating and hypertrophic zones of the growth plate. Dashed lines represent median (red dashes) and first and third quartiles (black dashes). p**** < 0.0001 (Kolmogorov–Smirnov test adjusted for multiple comparisons). n = 18 mice (2074 to 6359 cells per zone) (D) Major/longitudinal axis (red line) of cells in the proliferating zone of the growth plate. Scalebar: 5 µm. (E) Distribution of θ_cell,_ orientation angle with respect to the medial-lateral plane, in the resting, proliferating and hypertrophic zones of the middle and lateral regions of the growth plate. Dashed lines represent median (red dashes) and first and third quartiles (black dashes). p** = 0.0024, p*** = 0.00016, p**** < 0.0001 (Kolmogorov–Smirnov test adjusted for multiple comparisons) n = 18 mice (947 to 2737 cells per zone/region). (F) Percentage ciliation by growth plate region. Mean ± SEM. p*** = 0.0007 (paired t test). (G) Percentage ciliation in the different zones and regions of the growth plate. Mean ± SEM. p**** < 0.0001. All other comparisons are not statistically significant (two-way ANOVA). n = 18 mice (947 to 3622 cells per zone/region). (H) Primary cilia length in the different zones across the width of the growth plate. Mean ± SEM. p** = 0.0031, p*** = 0.0004 (one-way ANOVA) n = 18 mice (1028 to 3391 cilia per zone). Each data point is the average cilium length in the growth plate zone of one mouse. (I) Primary cilia length in the resting, proliferating and hypertrophic zones of the middle and lateral regions of the growth plate of a 6-week old Ift88^CTRL^ mouse. Mean ± SEM. p* = 0.0198. All other comparisons are not statistically significant (two-way ANOVA). n = 18 mice (466 to 1922 cilia per zone/region). Each data point is the average cilium length in the growth plate zone and region of one mouse. (J) Ki-67 staining in the growth plate of a 6 week-old Ift88^CTRL^ mouse. Scalebar: 50 µm.(K) Percentage of Ki-67 positive cells in the Ift88^CTRL^ Mean ± SEM (unpaired t test) n =3-19 mice.

Ciliation was 50% and 60% respectively between central and lateral regions (p<0.01). A statistically significant difference in % ciliation was present when comparing middle to lateral region (Figure 2G). Increased ciliation was most marked in the resting zone of the lateral growth plate (Figure 2H). Cilia length (Figure 2I) was largely between 1 and 4um. While no large differences were seen across zone or region, cilia lengths were statistically significantly longer in hypertrophic zone in middle region.

### Ciliation is reduced in peripheral, lateral hypertrophic regions, where IFT88 cKO phenotype is most prevalent

Our previous work described the disruptive, mechano-dependent, effects of IFT88 deletion in the articular cartilage[36] and growth plate [22]. Both implicated cilia in hypertrophy. Our new pipeline allowed us to quantify reductions in cell size, most notably in hypertrophic cells, with IFT88cKO (Figure 3A). These were also associated with changes to cell orientation (relative reductions towards outside of the limb) in the periphery of the limb in the resting and proliferative zones indicative of a shift away from horizontal (Figure 3B). Comparisons of ciliation revealed a reduction in cilia number in cKO in the periphery and hypertrophic zones, with minor effects on cilia length. As such this linked, in spatial manner, changes in IFT88cKO GP morphology, with changes to cells and cilia within these areas.

**Figure 3.**
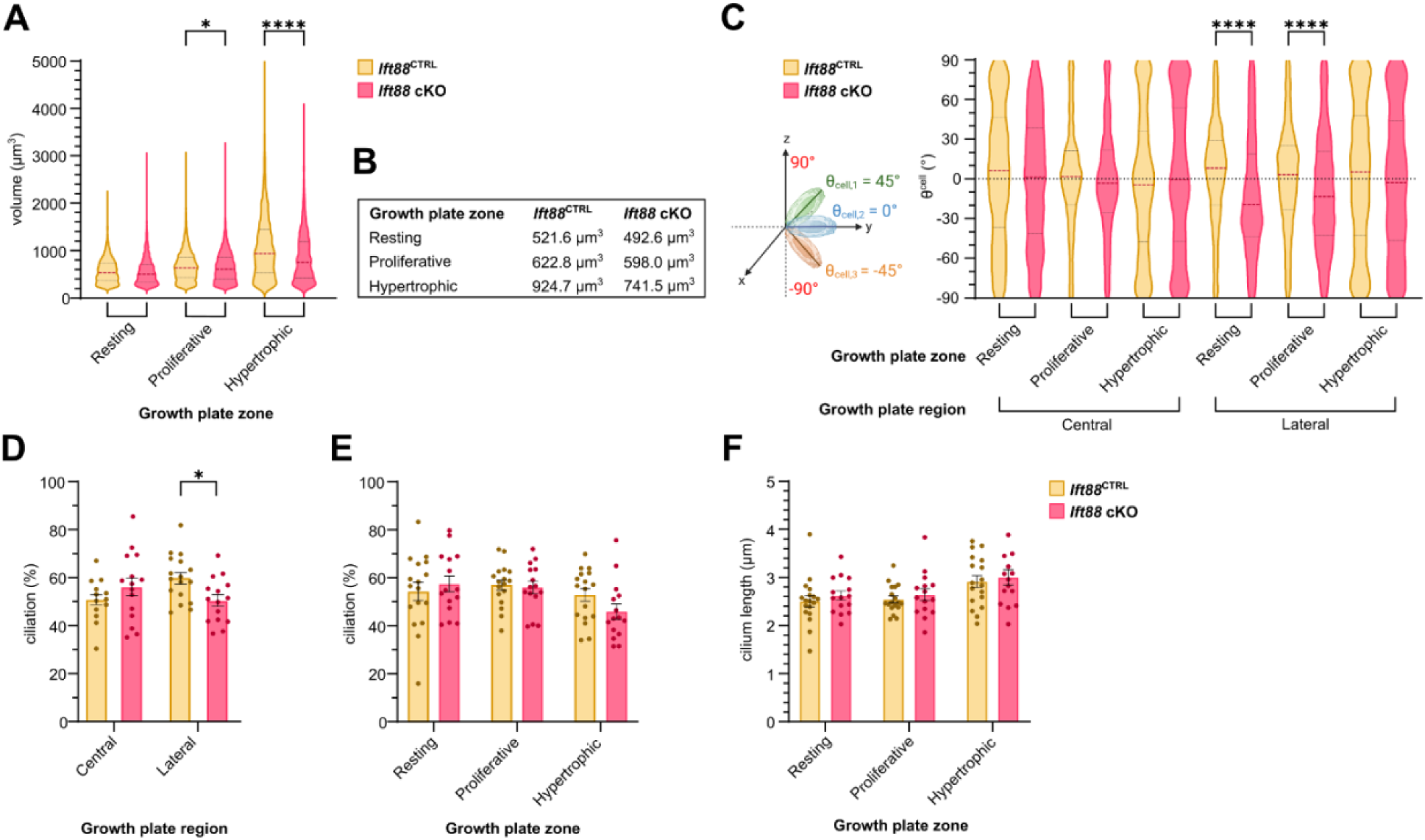
Cell size, orientation and ciliation in IFT88cKO. (A/B) Cell volume in the resting, proliferating and hypertrophic zones of the Ift88^CTRL^ and Ift88 cKO growth plates. Dashed lines represent median (longer dashes) and first and third quartiles (shorter dashes). Black asterisks – p* = 0.0161, p*** = 0.00013, p**** < 0.0001 (Kolmogorov–Smirnov test adjusted for multiple comparisons).n = 14-18 mice (585 to 3622 cells per zone/region). (B) Median cell volumes in the Ift88^CTRL^ and Ift88 cKO growth plates. (C) Distribution of θ_cell_ in the resting, proliferating and hypertrophic zones of the middle and lateral regions of the Ift88^CTRL^ and Ift88 cKO growth plates. Dashed lines represent median (longer dashes) and first and third quartiles (shorter dashes). p* =0.042, p** = 0.0018, p**** < 0.0001. (Kolmogorov–Smirnov test adjusted for multiple comparisons). n = 14-18 mice (585 to 3622 cells per zone/region).(D) Overall percentage ciliation (unpaired t test). (E) Percentage ciliation by growth plate zone. p* = 0.0291. All other comparisons are not statistically significant (two-way ANOVA). (F) Percentage ciliation by growth plate region. p* =0.0138 (two-way ANOVA). Mean ± SEM. n = 14-18 mice (301 to 1922 cilia per zone/region).

### Cell centrioles are non-preferentially positioned

First, we considered cell polarity, indicated by centriole pair (centrosome) position on cell. This is the site of ciliary assembly. Analysis revealed, across 14-18 mice assessing 1105 to 8204 centrioles per zone/region, that position was not preferential, in any axis, in any region or zone or through clonal column populations. For the centrioles, the two spherical coordinates Φ_centriole_ and θ_centriole_ represent the position of the centre of each centriole relative to the centre of the cell (Figure. 4A). Φ_centriole_ in the resting and proliferating zones showed broad distributions centred between 40° and 120°, suggesting centrioles were positioned mainly away from the proximal-distal positions. The resting zone of the middle region had a higher median than the two other zones. Hypertrophic centrioles presented a more spread-out distribution across all angle values. Many of the distributions were statistically different from each other, however the positions of centrioles they span were generally similar. The θ_centriole_ distribution is broad across all angles between 0 and 360° (Fig. 4E, F, G). There were slightly more centrioles at two modes around 50 and 300°. The median in the proliferating zone of the middle region was also higher than others. Though distributions were statistically significantly different from each other between certain zones and regions, the positioning of centrioles remained similar across the whole angle spectrum. Figures 4C-G show the same data of all centrioles across the growth plate. Figures 4D, G and H are rose and circular plot visuals demonstrating the general distribution of centriole positions across all spherical coordinates. On the cell surface, this translates as the centrioles being positioned at any point around the cell (Figure 5A). Appreciating that not all cells are ciliated, we asked whether there was a difference in the position of centrioles between cells that formed a cilium and those that did not. The position of the centrioles of ciliated cells and non-ciliated cells did not show any difference.

**Figure 4.**
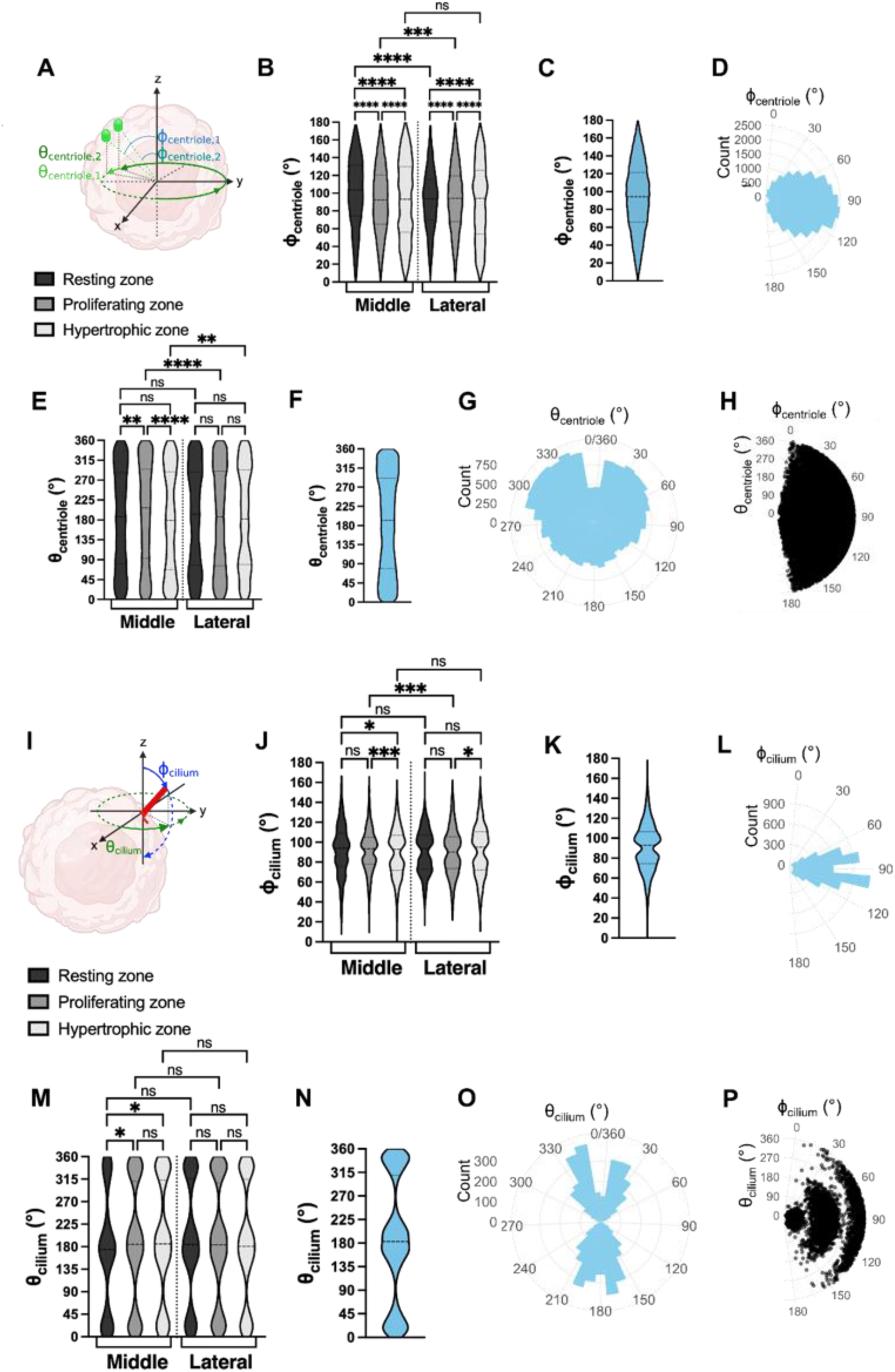
Centriole position and cilia axoneme orientation. (A) Spherical coordinates of centrioles. (B) Distribution of centriole Φ_centriole_ angles in the resting, proliferating and hypertrophic zones of the middle and growth plate. Dashed lines represent median (longer dashes) and first and third quartiles (shorter dashes). p***= 0.00024, p**** < 0.0001 (Kolmogorov–Smirnov test adjusted for multiple comparisons).n = 18 mice (1885 to 8204 centrioles per zone/region). (C) Distribution and (D) rose plot of Φ_centriole_ angles across the whole growth plate. n = 18 mice (25,740 centrioles). (E) Distribution of centriole θ_centriole_ angles in the resting, proliferating and hypertrophic zones of the middle and growth plate. Dashed lines represent median (longer dashes) and first and third quartiles (shorter dashes). p** = 0.0016 (resting middle vs proliferating middle), p** = 0.0023 (hypertrophic middle vs hypertrophic lateral), p**** < 0.0001.(Kolmogorov–Smirnov test adjusted for multiple comparisons). n = 18 mice (1885 to 8204 centrioles per zone/region). (F) Distribution and (G) rose plot of θ_centriole_ angles across the whole growth plate. n = 18 mice (25,740 centrioles). (H) Circular scatter plot of Φ_centriole_ and θ_centriole_ angles across the whole growth plate. n = 18 mice (25,740 centrioles). Primary cilia orientation is consistent throughout the 6-week old Ift88_fl/fl_ mouse growth plate. (I) Spherical coordinates of primary cilia. (J) Distribution of primary cilia Φ_cilium_ angles in the resting, proliferating and hypertrophic zones of the middle and lateral regions of the growth plate. Dashed lines represent median (longer dashes) and first and third quartiles (shorter dashes). p* = 0.022 (resting middle vs hypertrophic middle), p* = 0.016 (proliferating lateral vs hypertrophic lateral), p*** = 0.00082 (proliferating middle vs hypertrophic middle), p*** = 0.00082 (proliferating middle vs proliferating lateral) (Kolmogorov–Smirnov test adjusted for multiple comparisons). N = 18 mice (466 to 1922 cilia per zone/region) (K) Distribution and (L) rose plots and circular scatter plot of Φ_cilium_ angles across the whole growth plate. n = 18 mice (5802 cilia). (M) Distribution of primary cilia θ_cilium_ angles in the resting, proliferating and hypertrophic zones of the middle and lateral regions of the growth plate. Dashed lines represent median (longer dashes) and first and third quartiles (shorter dashes). p* = 0.047 (resting middle vs proliferating middle), p* = 0.047 (resting middle vs hypertrophic middle) (Kolmogorov–Smirnov test adjusted for multiple comparisons). n = 18 mice (466 to 1922 cilia per zone/region). (N) Distribution and (O) rose plots and circular scatter plot of θ_cilium_ angles across the whole growth plate. n = 18 mice (5802 cilia). (P) Circular scatter plot of Φ_cilium_ and θ_cilium_ angles across the whole growth plate. n = 18 mice (5802 cilia).

**Figure 5.**
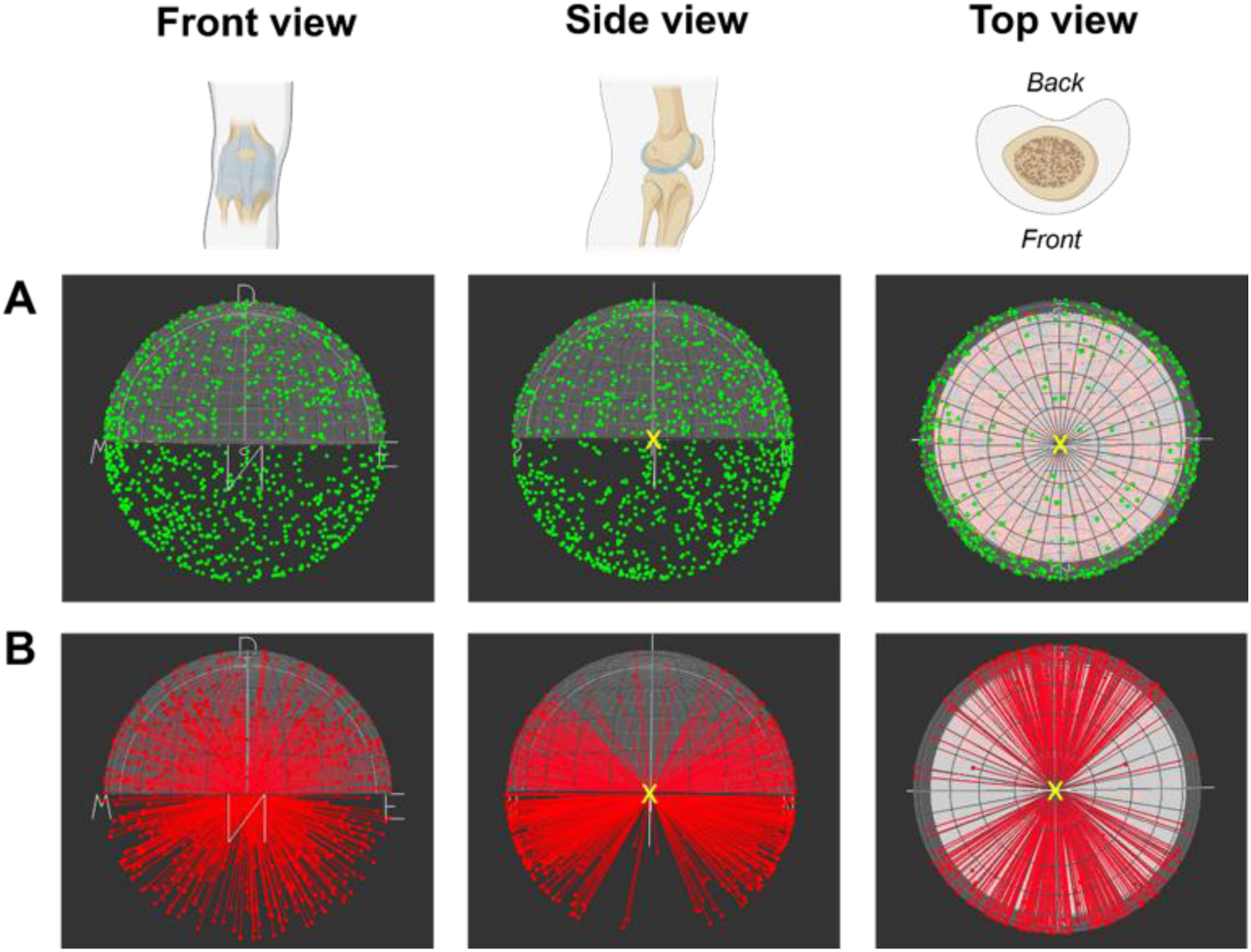
Centriole and primary cilia axoneme anatomical organization. Centriole (A) and primary cilia axoneme (B) organisation in the resting zone of a 6-week old Ift88^CTRL^ growth plate. For centrioles, the yellow cross denotes the centre of the cell with centriole position relative to this centre. For primary cilia, the yellow cross denotes where the cell would be with cilia pointing out. The geological software Stereonet was used for visualization.

**Figure 6.**
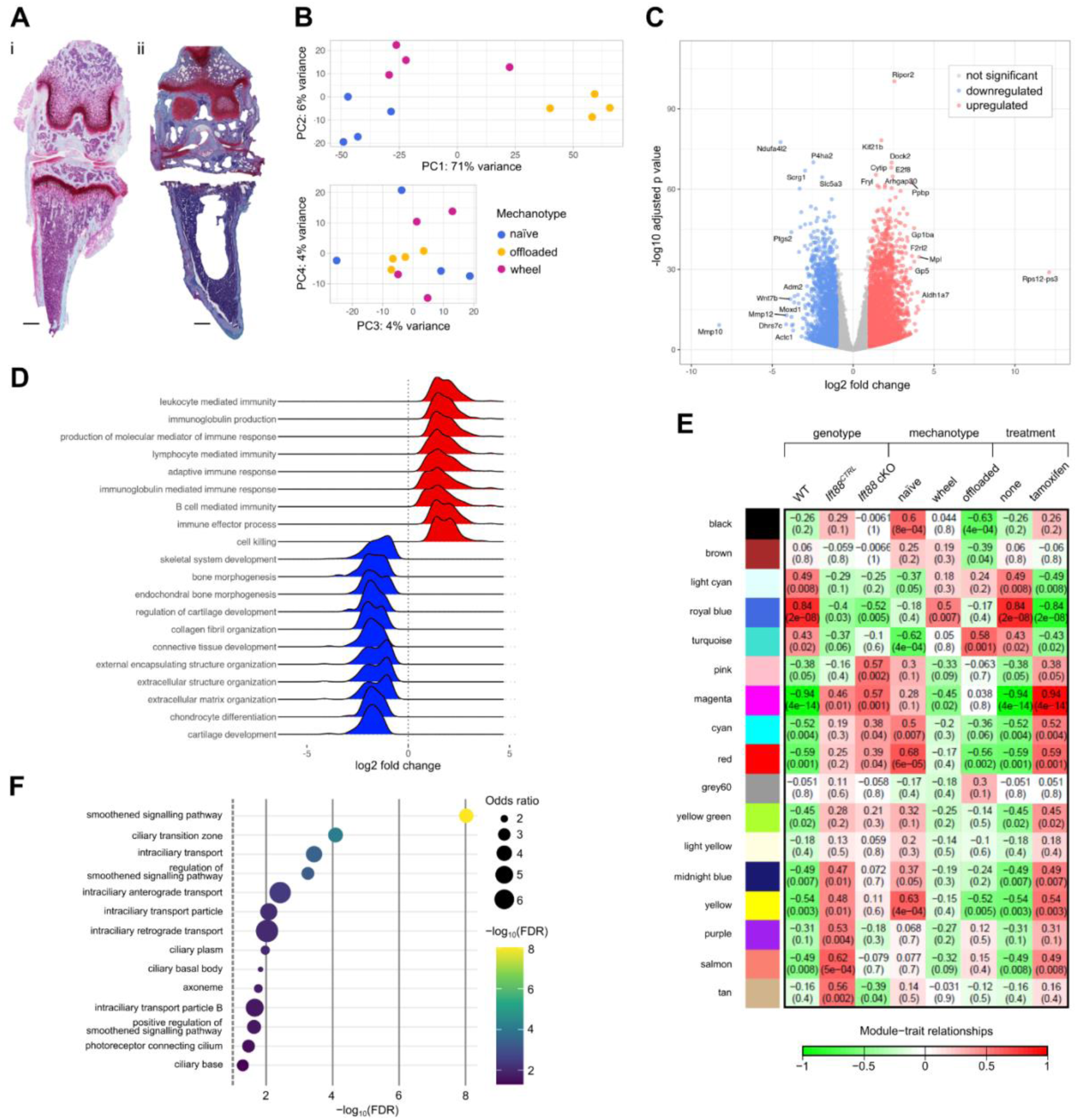
Transcriptional analysis of naïve and immobilised limb growth plate. (A) Histology (Safranin O/Fast green staining) before and after tibial growth plate removal. Scalebar: 500 µm. (B) PCA plots for wild type mice, naïve, immobilised and with wheel exercise. (C) Volcano Plot depicts differential gene expression analysis comparing naïve with immobilised. (D) Ridge plot depicts distribution of log2 fold changes for the genes found in corresponding top-most enriched GO terms. (E) Correlation analysis between module genes and sample traits. (F) Over-representation analysis of genes enriched in the black module selected ciliary associated GO terms shown (top terms shown in Table 1.).

### Primary cilia axoneme are preferentially orientated in the growth plate

Next we considered axoneme orientation. Strikingly, in contrast to centriole position, axonemal orientation was clearly not evenly distributed across all 3D angles. To measure 3D orientation of primary cilia, the spherical coordinates Φ_cilium_ and θ_cilium_ were measured across the different zones and regions of the growth plate. The two angles represent the direction the axoneme of the cilium is pointing in (Figure 4I). The Φ_cilium_ distribution was between 40° and 140° for all zones and regions of the growth plate, exhibiting a symmetric pattern centred around 0° (horizontal orientation) (Figure 4J). All areas had fewer cilia with Φ_cilium_ around 90°except the resting zone of the middle region. The other distributions had modes around 70° and 100°. This distribution suggests an inclination of primary cilia that is horizontal, but not perfectly horizontal (Φ_cilium_ = 90°). The proliferating zone cilia consistently had a different distribution of orientations compared with the hypertrophic zone in both regions. The proliferating cilia distributions were different between the two regions. Figure 4L/O represents the same data as Figure 4K and N in a rose plots which provides a visual perspective of all the angles across the whole growth plate. The θ_cilium_ angles were clustered around ∼30°, ∼180°and ∼350° (Fig. 4M). Visually, this corresponds to the cilia pointing forwards and back or anatomically dorsal and ventral (Figure 5B). The only difference in distribution was observed in the resting zone of the middle region compared with all the other regions and zones, where there were fewer cilia clustered around ∼350°. This analysis shows that primary cilia were non-randomly oriented in both elevation and planar direction, across thousands of cilia from multiple regions across 18 mice. There were few significant differences between the zones and regions of the growth plate, with the exception of the resting zone of the middle region which presents slight difference in terms of Φ_cilium_ and θ_cilium_ distribution.

Figure 5 gives a visual perspective of what this spherical coordinate distribution looks like in 3D, with cilia pointing primarily horizontally slightly up and down, and to the front and back. As such, whilst centriole position (green, A) is not anatomically preferentially positioned in 3D, ciliary axoneme orientation is (red, B). Critically, ciliary axonemes were not pointed parallel to limb or medially or laterally, but are orientated to the posterior (back) or anterior (front) of the limb usually at 45 degrees to limb axis.

### Broad effects of limb immobilisation on growth plate transcriptome include a ciliary signature

It has long been speculated that primary cilia position and/or orientation is mechanically regulated. Ambulatory loading regulates cellular organisation in the GP [52] and cilia are mechanosensors in other systems [17, 53]. In order to test this we exploited surgical immobilisation, used previously to reveal the mechano-dependence of IFT88 role in growth plate [22]. As a relative comparator only, alongside naïve relatively sedentary movement, we also used voluntary wheel exercise, previously shown to exacerbate the phenotype [22]. Later, we considered the contralateral limb, which receives normal or slight overloading as a comparative, and paired, control. Surgical immobilisation (off-loading) was conducted in one limb over a two-week period. After this the growth plate was collected for transcriptomic analysis by bulk RNA sequencing (Figure 6A). PCA analyses immediately revealed marked differences, particularly between naïve (normal ambulatory loading) and off-loaded (immobilised) groups (Figure 6B). A differential expression analysis revealed substantial (20% of analysed genes) transcriptional differences between wild-type naive and immobilised samples. Of the 4060 (statistically significant) differentially expressed genes (BH adjusted p value < 0.05), 2351 were upregulated and 1709 were downregulated. Top up-regulated genes in the immobilized condition include Ripor2(Rho-interacting in stereocilia), Dock2 (actin reorganization), and Re1n (extracellular matrix protein Reelin), while downregulated genes include Mitochondrial enzyme, Ndufa412, P4ha2, involved in collagen synthesis, and Col10a1, a marker of hypertrophy in the growth plate. Geneset enrichment analyses (GSEA) against the Gene Ontology Biological Processes database uncovered 1192 significant gene sets and notably revealed a marked downregulation in genes associated with skeletal differentiation and a strong upregulation of immune cell related terms (Figure 6D). We hypothesised that this upregulation of immune cell genes was associated with material changes to the spongiosa below the GP that resulted in increased representation of myeloid cells in the sample.

To further characterise the effect of mechanical status, versus genotype and experimental variables such as tamoxifen (controlled for in experimental comparisons) and caging, we applied weighted-correlation network analysis (WGCNA) to cluster genes showing similar expression patterns across all samples into “modules”. We then tested the association between the identified genes modules and sample characteristics (traits), enabling us to identify sets of genes that were associated with mechanical and/or genetic modifications but not with potentially confounding factors. As described in methods, caging and thus also therefore date of collection, largely overlapped with sample groups so Figure 6E shows experimental factors we are concentrated on; genotype and ‘mechanotype’ and the effects of tamoxifen. All modules are fully described in Supplementary Table 1 depicting top pathways and gene pathways in each.

Notably among the WGCNA gene modules, turquoise splits off first in the eigengene dendrogram (Supplementary Figure 5A) and expression is strongly associated with immobilisation (Supplementary Figure 5B). At >6000 genes it is the largest module. Top GO BP gene sets include protein localisation to chromosome, kinetochore and centromere and a strong signature related to DNA replication (Supplementary figure 5C). The other standout characteristic is a strong lymphocyte and adaptive immunity signature, thus turquoise represents a large component of up-regulated genes upon immobilisation (Figure 6D).

The next two largest modules are black and brown, both also associated with mechanotype (Figure 6E) but without this cell division/immune signature. Top GO BP terms for these modules include smoothened signalling and cartilage development respectively (Supplementary Figure 5C). Brown module also contains genes associated with non-motile or primary cilia and the black module contains genes with structural, functional and ciliary signalling (e.g smoothened, a component of hedgehog signalling [54]) genes (Figure 6F). A further examination of the black module indicates this signalling component, and its regulation, are joined by IFT transport genes including IFT88 itself. The effects of tamoxifen, whilst controlled for in these experiments as genotypes and mechanotypes both received tamoxifen, is the subject of a separate study.

**Supplementary Figure 5.**
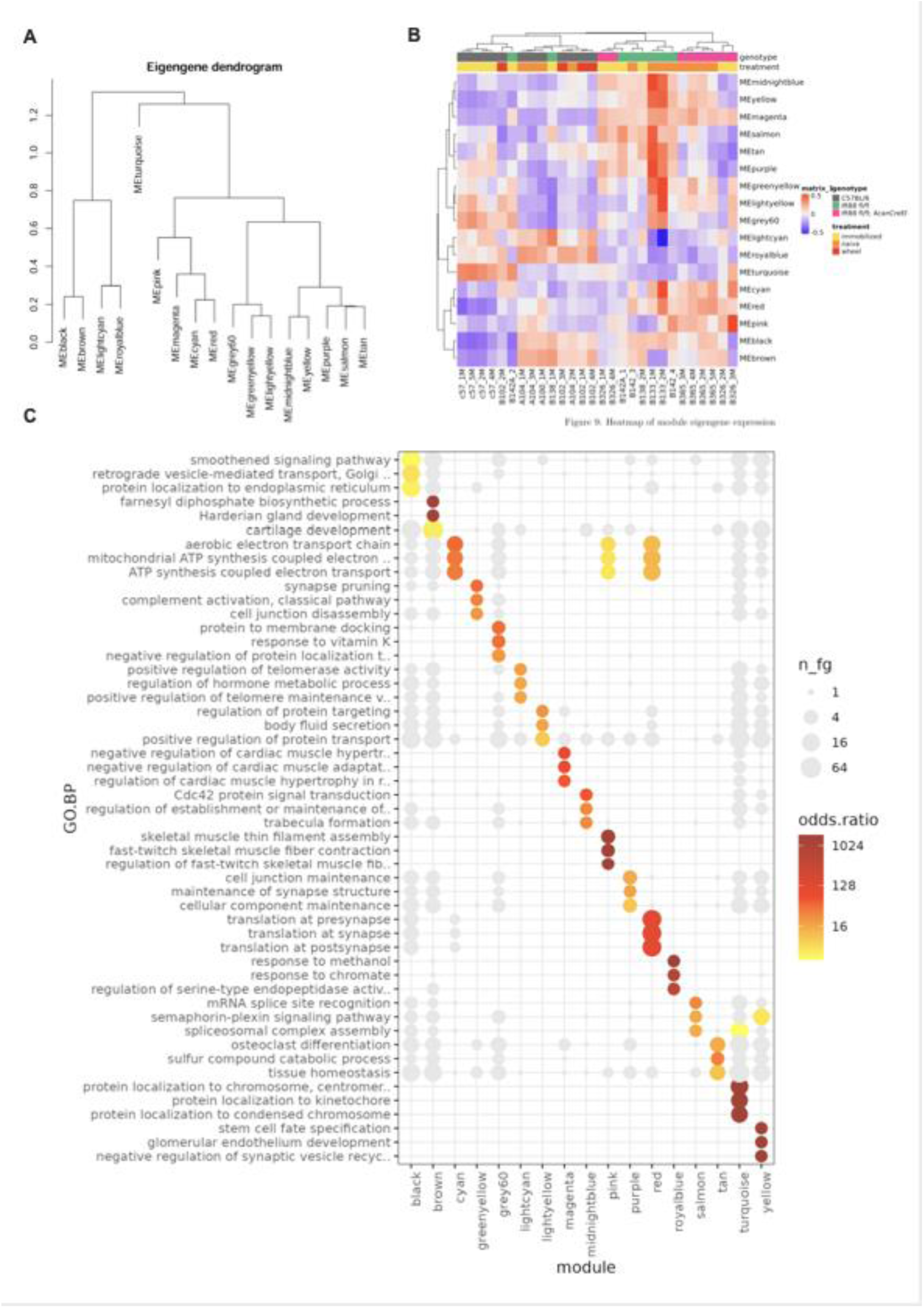
Full description of WGCNA Modules. (A) depicts the dendrogram as modules are split out. (B) Heatmap of module eigengene expression. (C) Top GO BP terms for each module, fuller descriptor in Supplementary Table 1.

The table lists the top (filtered by nominal p value) genesets by cluster.

**Table 1:**
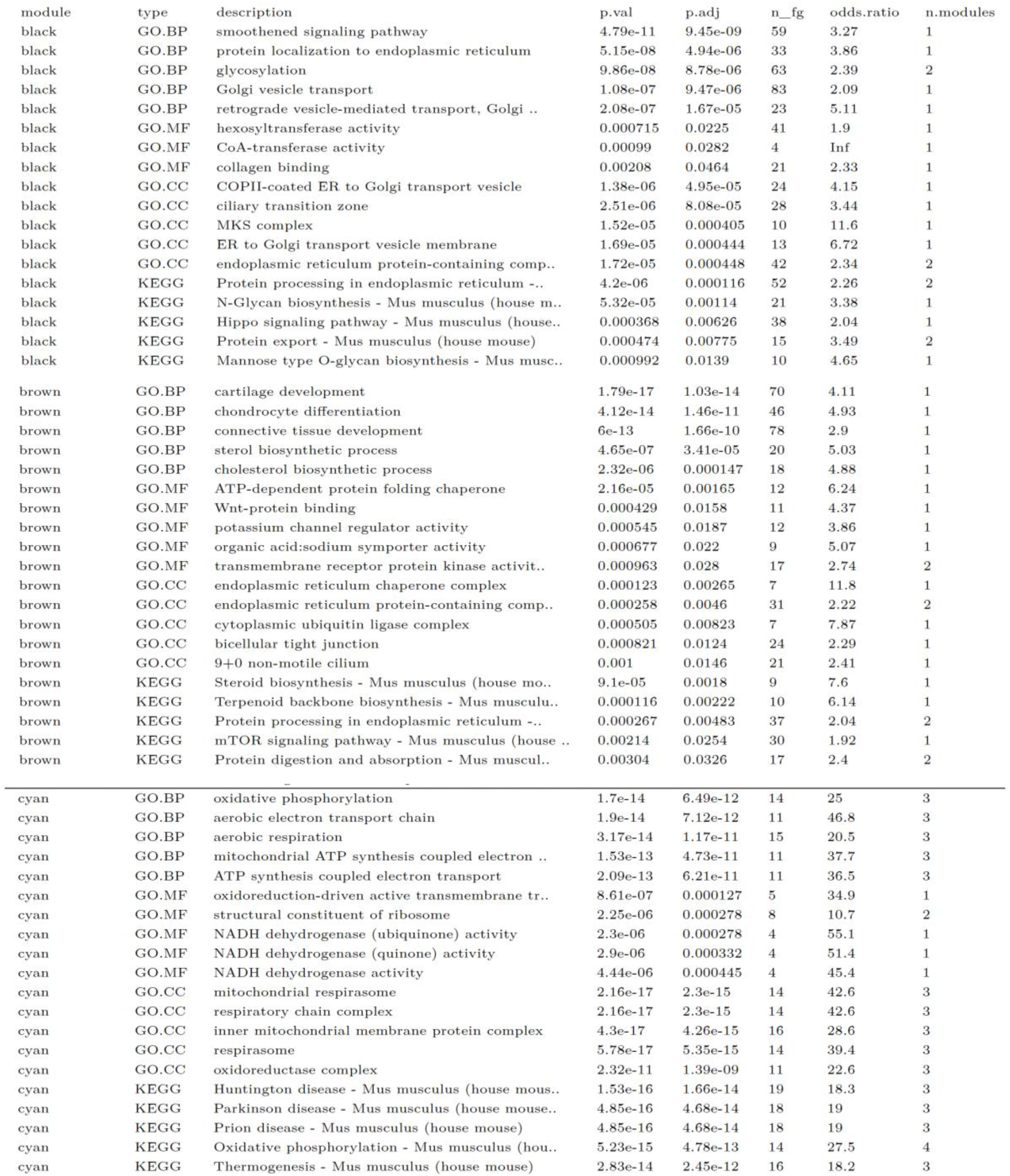
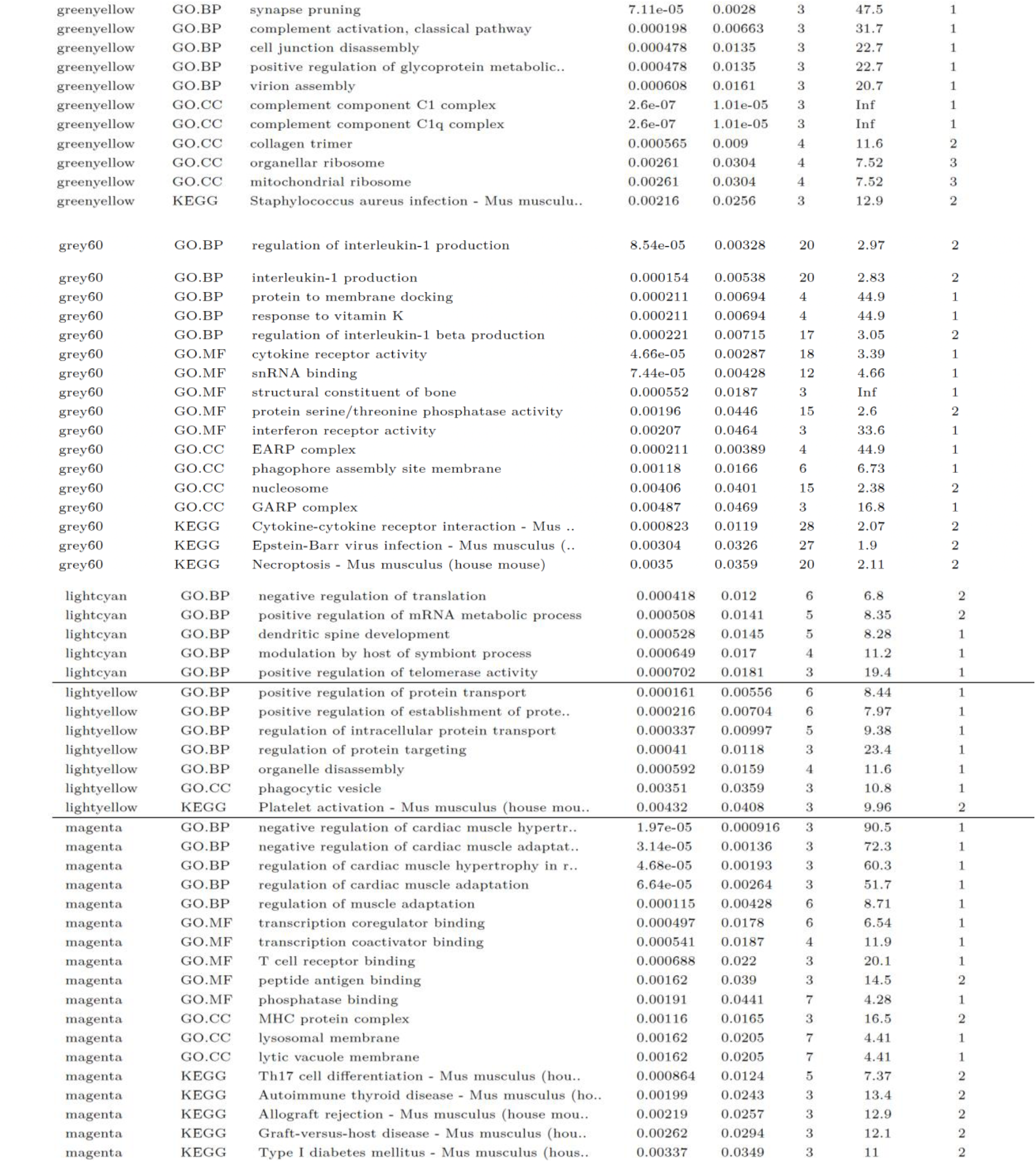
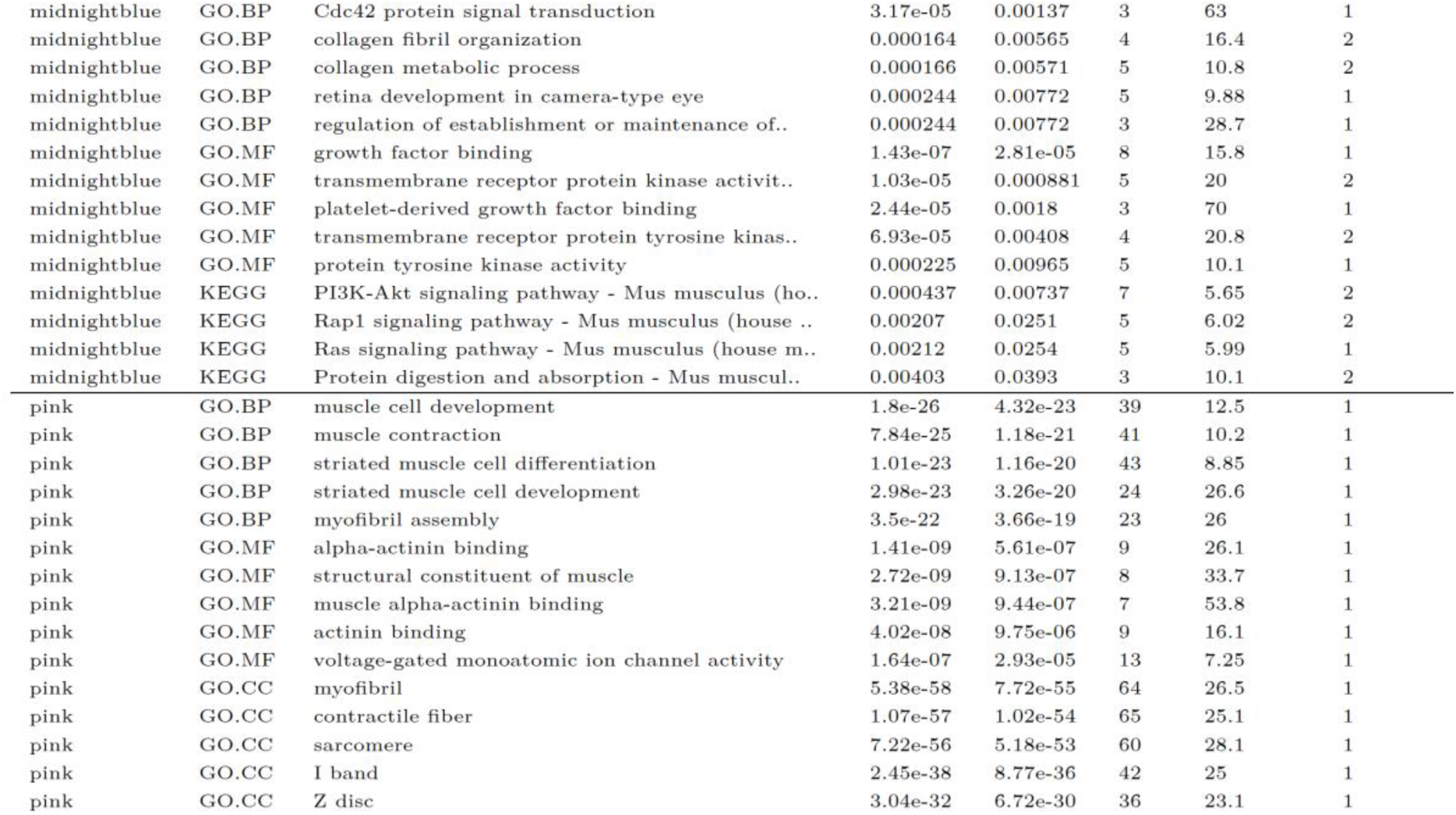
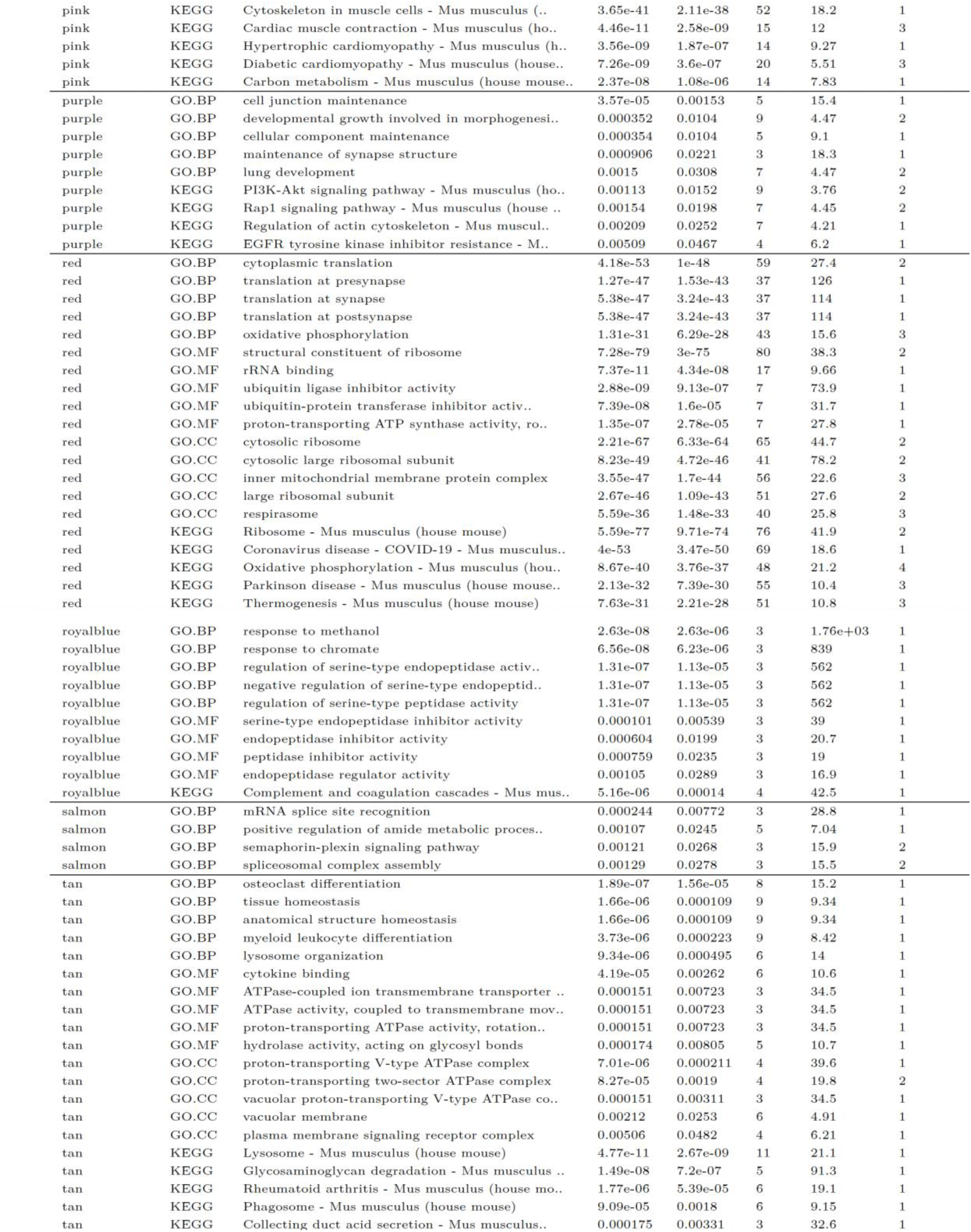
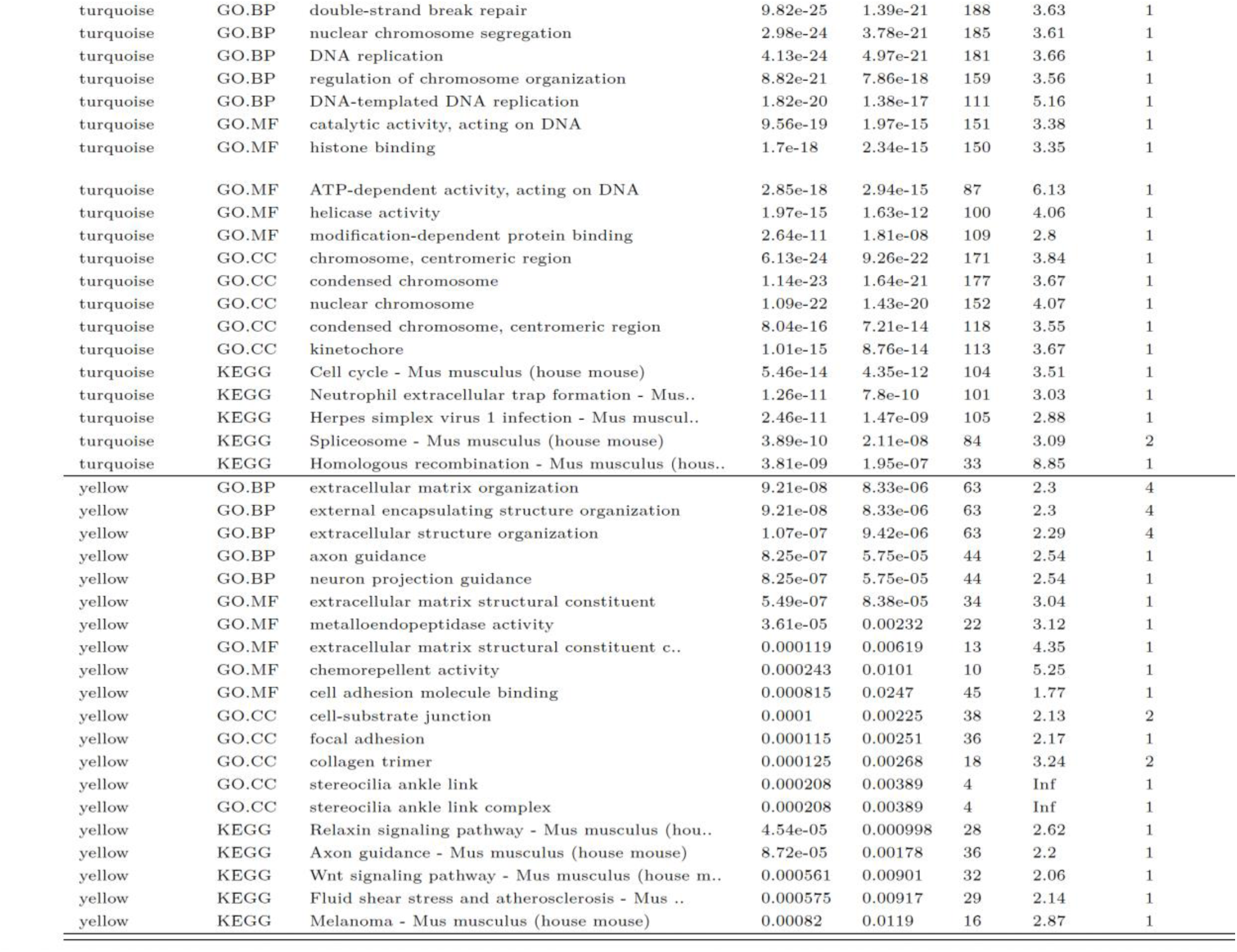
The top (lowest p-value) genesets found (uniquely) in each module.

**Supplementary Table 1**: *Full description of top genesets in each module*.

### Epiphyseal cellular morphology is altered upon limb immobilisation

Using the pipeline we next assessed cell size and orientation in immobilised and contralateral limbs. This identified that cell orientation changes in the resting zone, in a similar manner to changes seen with IFT88cKO. Immobilisation impairs the relative cell swelling (hypertrophy) in hypertrophic regions. These changes are associated with changes to GP length (Figure 8A-D).

**Figure 7.**
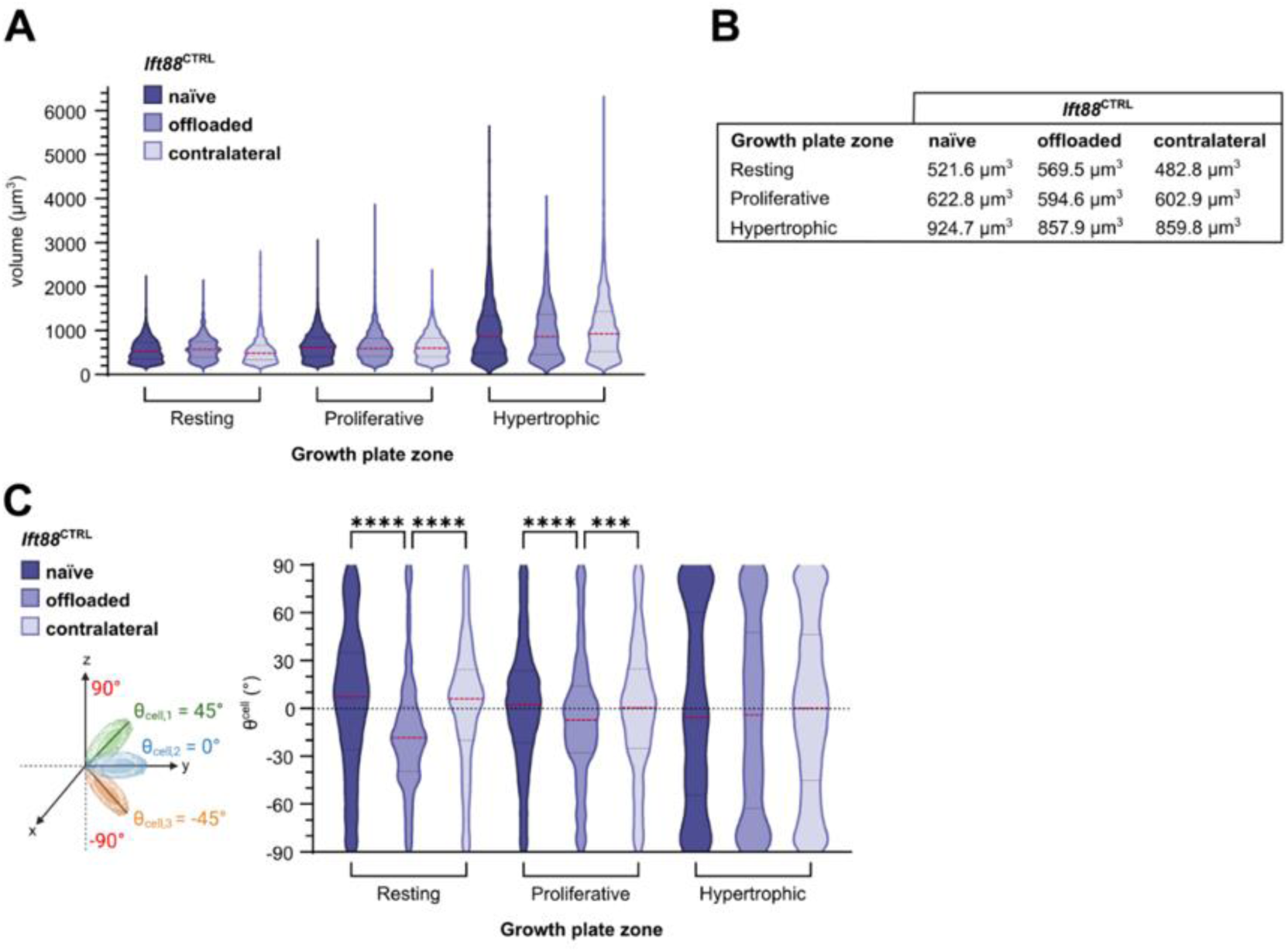
Cell volume and orientation upon immobilisation. (A) Cell volume in the resting, proliferating and hypertrophic zones of the control, offloaded and contralateral growth plates. Dashed lines represent median (longer dashes) and first and third quartiles (shorter dashes). p*** = 0.00024, p**** < 0.0001 (Kolmogorov–Smirnov test adjusted for multiple comparisons). n = 18 mice (2074 to 6349 cells per zone) (naive), n = 4 mice (332 to 1715 cells per zone) (offloaded, contralateral). (B) Median cell volumes in the control, offloaded and contralateral growth plates. (C) Distribution of θcell in the resting, proliferating and hypertrophic zones of the control, offloaded and contralateral growth plates. Dashed lines represent median (longer dashes) and first and third quartiles (shorter dashes). p* = 0.019, p**** < 0.0001. (Kolmogorov–Smirnov test adjusted for multiple comparisons). n = 18 mice (2074 to 6349 cells per zone) (naive), n = 4 mice (332 to 1715 cells per zone) (offloaded, contralateral).

**Figure 8.**
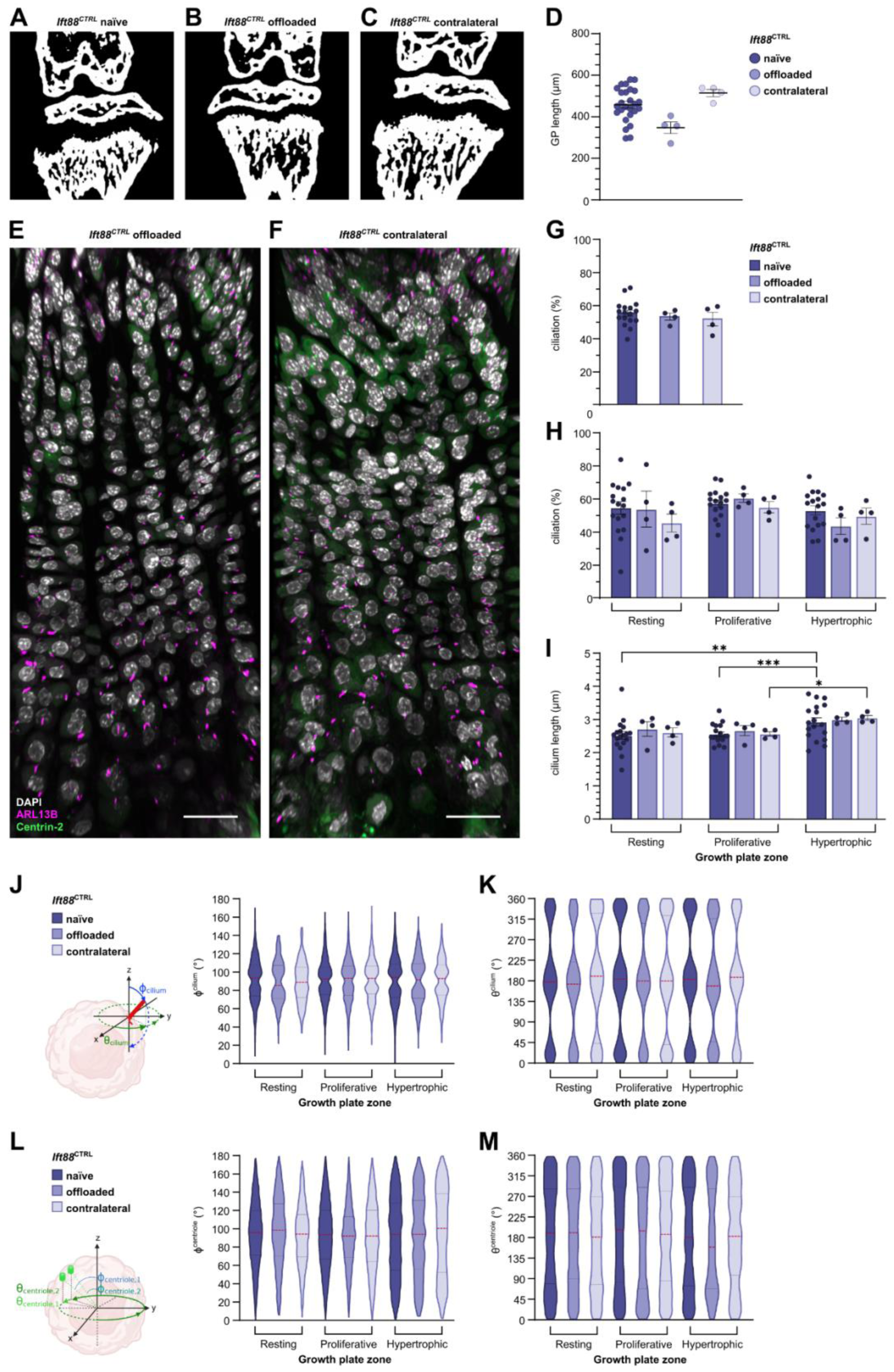
Effect of immobilization on primary cilia. Differences in growth plate length measured from thresholded microCT images of the naïve(A), Offloaded (B) and contralateral (C) Ift88^CTRL^ growth plates. (D) Thresholded growth plate lengths ofnaïve, offloaded and contralateral limbs. Mean ± SEM. p* = 0.0168, p** = 0.0023 (male) (one-way ANOVA) p = 0.005988 (female) (unpaired t test corrected for multiplicity).n = 4-18 mice. Primary cilia imaging in the middle of the growth plates offloaded (E) and contralateral (F) limbs. Scalebar: 30 μm. Percentage ciliation in the CTRL, offloaded and contralateral growth plates. (G) Overall percentage ciliation (one-way ANOVA). (H) Percentage ciliation by growth plate zone. All comparisons were not statistically significant (two-way ANOVA). Mean ± SEM. n = 14-18 mice (1660 to 6349 cells per zone) (Ift88^CTRL^, n = 4 mice (332 to 1715 cells per zone) (offloaded, contralateral). (I) Cilia length Mean ± SEM. p* = 0.0240 (proliferating vs hypertrophic Ift88^CTRL^ contralateral), p** = 0.0031 (resting vs hypertrophic Ift88^CTRL^), p*** = 0.0004 (proliferating vs hypertrophic Ift88^CTRL^). All other comparisons were not statistically significant (two-way ANOVA). n = 14-18 mice (778 to 3299 cilia per zone), n = 4 mice (128 to 901 cilia per zone) (offloaded, contralateral). (J) Differences in primary cilia orientation between the 6-week old CTRL, offloaded and contralateral growth plates. Distribution of primary cilia Φ_cilium_ angles in the resting, proliferating and hypertrophic zones across the growth plates. Dashed lines represent median (longer dashes) and first and third quartiles (shorter dashes). n = 14-18 mice (778 to 3299 cilia per zone) (Ift88^CTRL^), n = 4 mice (128 to 901 cilia per zone) (offloaded, contralateral). (K) Distribution of primary cilia θ_cilium_ angles in the resting, proliferating and hypertrophic zones across the growth plate. Dashed lines represent median (longer dashes) and first and third quartiles (shorter dashes) n = 14-18 mice (778 to 3299 cilia per zone) (ctrl), n = 4 mice (128 to 901 cilia per zone) (offloaded, contralateral). (L) Distribution of centriole Φ_centriole_ angles in the resting, proliferating and hypertrophic zones across the growth plates. Dashed lines represent median (longer dashes) and first and third quartiles (shorter dashes). n = 14-18 mice (2518 to 14,587 centrioles per zone) (Ift88^CTRL^), n = 4 mice (950 to 4611 centrioles per zone) (offloaded, contralateral). (M) Distribution of centriole θ_centriole_ angles in the resting, proliferating and hypertrophic zones across the growth plate. Dashed lines represent median (longer dashes) and first and third quartiles (shorter dashes) n = 14-18 mice (2518 to 14,587 centrioles per zone) (Ift88^CTRL^), n = 4 mice (950 to 4611 centrioles per zone) (offloaded, contralateral).

### Primary cilia structure and orientation is not regulated by ambulatory loading

We next used 2-week limb immobilisation, due to surgical intervention [55], to test the hypothesis that ambulatory loading orientates dynamic ciliary axoneme orientation. During 2 weeks, cells within the GP will divide multiple times and thus disassemble and reassemble cilia, in many cases having spun around within column [25], and certainly having moved into or created new matrix environments. Off-loading, through immobilisation results in a narrowing of the GP between 4 and 6 weeks of age whilst the contralateral length is comparable to control mice (Figure 8A-D). Due to only 4 mice being analysed for these experiments, analysis was performed across the growth plate (medial and lateral regions averaged) and presented by zone. The growth plates of the offloaded and contralateral limbs appeared ciliated with cells arranged in columns (Figure8 E/F). Percentage ciliation, primary cilia length and orientation were next measured to determine whether the absence of mechanical loading affected the axoneme of the primary cilium. The overall growth plate percentage ciliation in the offloaded and contralateral were the same as control, around 53% (Figure 8G). The percentage ciliation was also measured in the different zones to assess any differences at this level (Figure 8H). The offloaded growth plate had lower percentage ciliation in the hypertrophic zone compared with the resting and proliferating zones. The zones of the contralateral growth plate all had similar percentages of ciliation. The primary cilia lengths in the offloaded and contralateral growth plate followed a similar pattern to controls. The primary cilia were longer in the hypertrophic zone with a mean around 3 μm compared to ∼2.6 μm in the resting and proliferating zones (Figure 8I).

To determine whether absence of loading, or increased loading, changes the direction the axoneme of primary cilia point in chondrocytes, the spherical coordinates of the offloaded and contralateral growth plate cilia was measured. Loading in the limbs comes mainly from ambulatory load, therefore is directional in the proximal-distal direction. The absence of this directed load could be re-directing axoneme orientation during re-assembly. The offloaded and contralateral growth plates generally had primary cilia orientations that were similar to controls (naïve and contralateral) in terms of Φ_cilium_ and θ_cilium_ (Figure 8). The θ_cilium_ distribution was very similar across all conditions. This indicated that whilst mechanical loading influences dynamic transitions within the GP, ciliary axonemal orientation is not dynamically regulated in conditions of changes to mechanical loading over periods of weeks.

## Discussion

Our principal aim was to explore further into previous phenotypes, where the apparent effects of ciliary IFT88 deletion, were more pronounced in the GP periphery. We hypothesised that expanded hypertrophic regions were the result of hypertrophic to bone transitions being slowed, due to a role for cilia in chondro-osseous transitions. As such we set out to measure cilia prevalence in populations of GP chondrocytes comparing between the zones (depth) of the GP and across the width of the limb (central vs lateral periphery regions). The methods are previously described [6], but here we validate this, using predictable relationships in terms of cellular and tissue morphology, and exploit it to report our biological findings from 6-week old mice epiphyses. These data form a 3D description of cilia at organ level, allowing us to consider how primary cilia act at this scale, across collective cell populations. Combined with transcriptomic analyses, the data strengthen mechanistic links between the primary cilium and mechano-dependent regulation of stem cell and hypertrophic biology in the growing limb.

The methodology enables cell, ciliary axoneme and centriole detection at high resolution, enabling us to build an understanding of positions and orientations. With super resolution microscopy and Z resolution corrections, cilia axoneme orientation can be confidently mapped in 3D, using the spherical coordinates system and leveraging endogenous fluorescent markers and automated image processing [6]. This was done across tens of thousands of cells from 18 different animals growth plates. Cell volumes presented here are in-line with that of literature [56], but likely an underestimate, particularly for hypertrophic cells, as the GFP signal background used does not fill entire cell. Cell orientation is predictably, but measurably, altered through zones, as seen by a relative horizonal alignment in the vertical stacks of the proliferative zone. This reveals another difference between the middle and periphery of the growth plate. Central, resting zone, cells are more randomly orientated than peripheral (lateral) cells, potentially due to differentials in mechanical stresses [41, 57]. We show later that mechanical forces associate with ambulation can regulate this. The shape of the growth plate varies through its width, in line with the shape of SOC above. How this curvature affects cells within, the ECM, stresses and strains experienced, and orientation cues remains unknown. Proliferating cells have been previously described as the most organised in the growth plate, with the major axis of the cell paralleling the medial-lateral plane [58], [25, 38]. Hypertrophic cells lose the flattened, aligned, structure, as they expand in volume and prepare for trans differentiation or apoptosis [59]. The cellular column organisation in the growth plate is achieved through a coordinated mechanism of cell division and rotation. Following division in a plane parallel to the longitudinal axis of bone growth, newly produced daughter cells pivot around each other, stacking to form a column along the tissue’s elongation axis. The planar cell polarity pathway is essential for this coordinated behaviour [60]. However, despite this we find no preferential position of centrioles, often a proxy for polarity. ECM stiffness, mechanical cues and compressive forces are all implicated in promoting the alignment of columns along the longitudinal bone axis [52, 61, 62]. Notably, recent studies reveal variation during development. In embryos, column formation via full rotation of the division plane is less frequent, resulting in more clusters or irregular stacks, whereas in post-natal growth plates, complex columns –both ordered and somewhat disordered – are predominant [25]. This suggests the process is dynamic and regulated differently over time and across anatomical locations.

Cilia number is different between middle and periphery. Cilia prevalence is around 50% in the center of the limb, rising to 60% in the lateral periphery, a difference predominantly underpinned by increased prevalence in the resting zones comparatively between center and periphery (40% vs 60%). This may also relate to shape and strain changes, likely therefore exaggerated in the stem cell niche closest to SOC [23]. Quantifying cilia prevalence offers valuable information about the physiological relevance of primary cilia in a given tissue and here may be indicative of different stem cell populations in center and periphery. The ciliation percentages measured here are in the same range as previous reports, though measured at different ages and with slight differences in ratios of zones [22, 63, 64]. The method used here provides a more accurate counting method based on detection of cilia axoneme, compared with manual counting or immunohistochemistry (IHC) staining detection, and its high throughput allows for more regional detail. The differences in mechanical strains and micro-environments between the middle and lateral region highlighted above may explain the higher ciliation in the lateral region. The higher mechanical stresses experienced by the lateral region may justify its requirements for more cilia to integrate mechanical signals [41]. Modelling indicates that at the edges of the growth plate, chondrocytes are subject to higher transverse and shear strains [57]. These strains can trigger adaptive cellular responses, potentially upregulating ciliation as part of the tissue’s mechanosensory response. Our later experiments test whether differential mechanical load triggers such changes in ciliation in the growth plate. It could be expected that the proliferating region has the lowest ciliation, as cilia form when cells exit the cell cycle [65]. However, in tissue sections the cell division event in the proliferation zone where cells are side by side before rotating into columns is very rarely captured. It probably occurs very fast, and the cell state observed in sections is that of cells between cell divisions, thus able to ciliate. Ciliation at 50 and 60% respectively between middle and periphery is set against a total growth plate ki67% which ranges from 20-60% and qualitatively is diminished in the periphery. Ki67% reduces with age with no such effect on ciliation suggesting cell cycle status is not the sole regulator. As such, it remains unclear the relative roles mechanics, and the cell cycle, have to ciliation.

Primary cilia axonemal length, associated in many contexts with changes in function and chondrocyte biology [38, 58, 66–72], measurements reveal hypertrophic cells have the longest cilia. These cells appeared most influenced in models of cilia disruption [22]. This is in contrast with the shortest cilia measured in the hypertrophic zone of the humerus elbow in embryos [64]. This detection was based on acetylated α-tubulin, whereas here ARL13B signal is detected. As ARL13B is localised to the ciliary membrane, it can indicate the full length of the ciliary projection, including any membrane extensions beyond the axonemal core. Acetylated α-tubulin marks the microtubule-based structural core within the cilium, which may not reach the absolute tip of the ciliary membrane in every instance. Cilia length measurements based on acetylated α-tubulin have been shown to be different to measurements based on ARL13B [73–75]. Acetylated α-tubulin can also label non-ciliary microtubules, potentially leading to false positives, whereas ARL13B is more specific for primary cilia [76]. In addition, the function of primary cilia may differ between embryonic and postnatal stages, reflecting the distinct tissue architecture and extracellular matrix cues present at each stage.

Analysis of orientation and position reveals there is no preferential position of centrioles within the growth plate, at 6 weeks of age. This, like column formation, might be different at this time point to earlier and does not wholly speak to polarity, but does indicate that cilia position is not polarized. In contrast, cilia axoneme orientation is highly ordered with a preference across the population to non-random. In contrast to previous findings, there is preferential posterior and anterior, dorsal-ventral, orientation, 45 degrees to limb horizontal axis. This orientation is not altered in conditions of sustained, 2 weeks, immobilization. As discussed later, this removal of ambulatory loading affects cellular organization and, from previous publications [22, 43] and our own work, we know profoundly affects cell and matrix behavior in cartilage of the epiphysis and surrounding bone. Quite why cilia axoneme are orientated this way with respect to the limb remains unclear. Collectively cells will have a range of angles. Orientations do not relate apparently to axial vs radial stiffness or tissue anisotrophy, as they do not alter with region or zone unlike prevalence and length. To us it suggests a cell intrinsic mechanism perhaps shared with cilia in other tissues. Oscillatory orientations controlled by circadian rhythms have been observed in the brain [5].

The analysis in cKO mice, where IFT88 is conditionally ablated under the control of tamoxifen using an aggrecan cre driver [22] reveals that there are changes to cell size, notably reductions in hypertrophy, and changes to horizontal orientations of cells in resting zone and proliferative zones, these are in line with previous observations in other ciliary KO models [27]. The greatest changes to ciliation are in the periphery, in the hypertrophic regions but also in the stem cell zone (Figure 3). This means that in addition to the effects being demonstrative of a role in dealing with greater strains in the periphery, the effects in cKO are likely to be due to a greater impact on cilia in these regions. As previously reported [22], the Cre does not drive preferentially in any regions, but may drive preferentially in zones. Nevertheless, these data again link cilia axoneme to the IFT88cKO phenotype. It was logical to next explore how mechanical forces themselves influence cilia, as their assembly is mechanosensitive [68, 77].

Surgical immobilization allows 2 weeks of removal of forces through the growth plate associated with ambulatory loading. Morphological analyses have previously highlighted how such immobilization affects the growth plate [52], and how immobilization can effectively protect against disruption with loss of cilia [22]. RNAseq analysis identified large-scale transcriptional changes with approximately 20% of genes analyzed differentially expressed. Interestingly, immobilization was also associated with changes to the horizontal plane organization of cells in the GP, again in the resting zone, in line with previous observations [52] and to hypertrophy, again linking ciliary effects to changes in mechanobiology. Within the gene signatures identified, pathways relating to extracellular matrix regulation were enriched in the down-regulated genes and many apparently immune-associated genes increased. The increase in immune genes we ascribe to collection of softer, bone including marrow, due to effects of immobilization on calcification. To help deconvolute these complex changes in gene expression, we applied WGCNA to discover modules of genes that co-varied across experimental conditions. Within these, a strong ciliary signature emerged leading us to hypothesize that immobilization and ambulatory loading, regulates cilia themselves (IFT88 is regulated) and their activity in form of, for example, hedgehog (Hh) signalling. Hh signaling is central to EO, even in the adolescent growth plate [35, 78]. The removal of ambulatory loading in immobilization experiments indicates loss of normal movements for 2 weeks whilst influential to the GP, is not influential to cilia prevalence. This striking as ki67 staining indicates over 2 weeks these GP cells will have disassembled and re-assembled cilia after cell division at least once, but that axonemal orientation is unaffected. Immobilization, and the change of loading in the contralateral limb, does result in clear changes in growth plate length, which we identify in other studies to be associated with shifts in cell populations (increased in contralateral and reduced in immobilization). These parallel studies are describing the wide-ranging effects on limb biology in order to understand the role of cilia and mechanics in this context. Here, we describe that immobilization changes cell sizes and apparent changes in cell orientation, so it is surprising that cilia orientation is resilient to this. We think this implies an internal cell intrinsic mechanism or program drives orientation, given to the cell early or inherent to cells in the growth plate, without requirement for ambulatory loading.

To summarise, accurate imaging, and analysis, of cells and cilia across populations and regions, and powered over multiple animals, reveals patterns of ciliation. There is increased cilia prevalence in the limb periphery, especially in the resting zone which harbors stem cell populations. The periphery is where the phenotypic effects of IFT88 knock out are the greatest, and associated reductions in cilia number are the greatest, linking IFT88 effects directly to primary cilia. Analysis of position and orientation reveals a conserved and resilient orientation of the axoneme that is independent to cilia position/centriole position. The effects of immobilization on growth plate biology are wide-ranging and include effects on ciliary signaling. However, preferential orientation of cilia is not governed, it seems, by ambulatory limb loading. These data imply to us, an intrinsic mechanism or sustained role for matrix order. Cilia orientation is non-random, independent of centriole position and whilst cilia and ciliary signaling are central to how limb growth is mechano-regulated, why cilia are orientated this way remains unclear. Perhaps this allows collective decision making, something we are exploring now we have the *in vivo* ‘ground truth’ of ciliary mapping. Perhaps, the most important point is that whilst cilia are assembled from any position, axoneme are never parallel to forces through the limb (cranial-caudal, Z) or parallel to the medial-lateral plane (left to right, X) but can be parallel to the dorsal ventral plane (front to back, Y) (Figure 9). Our interpretation of this is that being Z or X parallel, would disable the ability to sense a gradient of force in say the X direction or a biological cue such as a growth factor gradient in the Z direction (Figure 9B). Interpreting gradients/signals in both directions is integral to morphogenetic processes, and we propose cilia orientation is critical.

**Figure 9:**
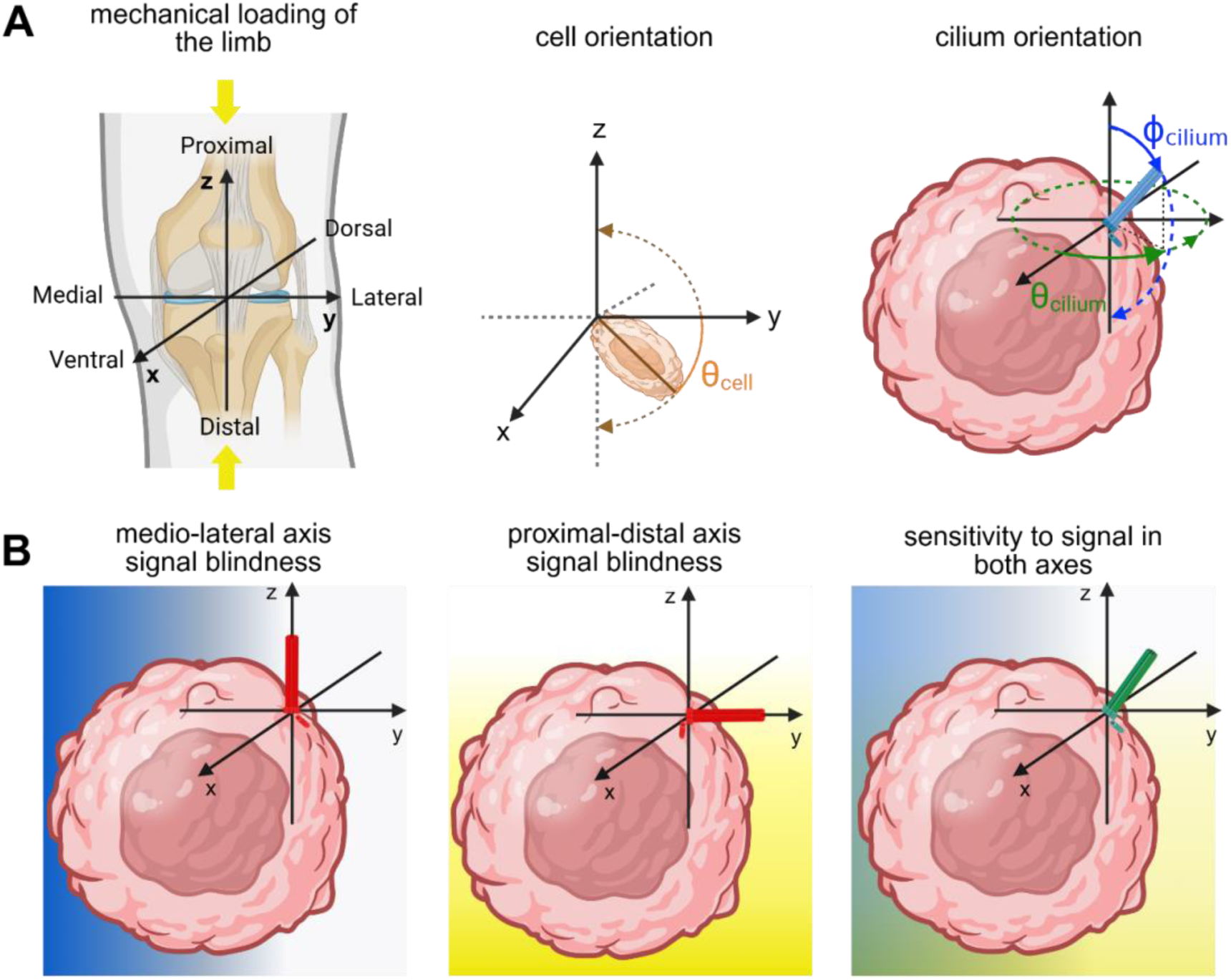
Schematic for conceptualization of ciliary axoneme orientation to avoid single X/Y/Z signal ‘blindness’. (A). Physical compressive forces are in the proximal-distal (cranial-caudal) Z direction, cells within GP may experience signals or gradients in medial-lateral or ventral-dorsal (Y or X) direction and Z direction. (B). Cilia are never orientated vertically or medial-lateral horizontal, an orientation that is not parallel to any single axis, enables integration of multiple signals and avoids blindness in any axis.

## Acknowledgement

The authors would like to acknowledge that without the transgenic line used previously [42] and now here, this study would not have been possible. As such great thanks go to Dr Fiona Bangs and the late Kathryn V Anderson. The work was hugely supported by those within the KIR; in the BSU notably Albertino Bonifacio, and by histology team; Ida Parisi, Oyindamola Ajisegiri and Rhiannon Cook. The authors also acknowledge the KIR Zeiss microscopy unit and Ka Long Ko for help with image analysis and the wider support of the Arthritis UK Centre for OA Pathogenesis. Schema throughout paper were created in Biorender.

